# Alveoli form directly by budding led by a single epithelial cell

**DOI:** 10.1101/2021.12.25.474174

**Authors:** Astrid Gillich, Krystal R. St. Julien, Douglas G. Brownfield, Kyle J. Travaglini, Ross J. Metzger, Mark A. Krasnow

## Abstract

Oxygen passes along the ramifying branches of the lung’s bronchial tree and enters the blood through millions of tiny, thin-walled gas exchange sacs called alveoli. Classical histological studies have suggested that alveoli arise late in development by a septation process that subdivides large air sacs into smaller compartments. Although a critical role has been proposed for contractile myofibroblasts, the mechanism of alveolar patterning and morphogenesis is not well understood. Here we present the three-dimensional cellular structure of alveoli, and show using single-cell labeling and deep imaging that an alveolus in the mouse lung is composed of just 2 epithelial cells and a total of a dozen cells of 7 different types, each with a remarkable, distinctive structure. By mapping alveolar development at cellular resolution at a specific position in the branch lineage, we find that alveoli form surprisingly early by direct budding of epithelial cells out from the airway stalk between enwrapping smooth muscle cells that rearrange into a ring of 3-5 myofibroblasts at the alveolar base. These alveolar entrance myofibroblasts are anatomically and developmentally distinct from myofibroblasts that form the thin fiber partitions of alveolar complexes (‘partitioning’ myofibroblasts). The nascent alveolar bud is led by a single alveolar type 2 (AT2) cell following selection from epithelial progenitors; a lateral inhibitory signal transduced by Notch ensures selection of only one cell so its trailing neighbor acquires AT1 fate and flattens into the cup-shaped wall of the alveolus. Our analysis suggests an elegant new model of alveolar patterning and formation that provides the foundation for understanding the cellular and molecular basis of alveolar diseases and regeneration.

**One Sentence Summary:** We report a direct budding mechanism of alveolar development distinct from the classical model of subdivision (‘septation’) of large air sacs.

## Introduction

The respiratory surface of the mammalian lung is formed by millions of regularly spaced, densely packed, thin-walled air sacs called alveoli arranged as single units or in small groups along the tubular walls or ends of airways (*1, 2*). Air passes along the airways into alveoli, where oxygen diffuses across the alveolar wall to reach circulating red blood cells, which distribute it throughout the body (*3*). Classic studies of lung structure revealed the geometry and architecture of respiratory airways (*1, 2, 4–8*) and provided stereological estimates of alveolar number (2-4 million in mice), size (50 µm mean diameter), and surface area (6 x 10^3^ µm^2^) (*9–12*). However, the extreme thinness of alveolar cells and dense packing of alveoli make it difficult to count individual cells and define their structures and boundaries on histological sections. Electron microscopy and serial section reconstruction resolved the fine structure of the air-blood barrier and the organization of the alveolar epithelium (*13–18*), but the techniques are laborious and have not been used to systematically define the morphologies, numbers and arrangement of the different cell types in an alveolus. Hence the three-dimensional cellular structure of alveoli has not been resolved.

There is an urgent need to define the cellular structure and formation of alveoli since they are the sites of major, life-threatening lung diseases. These include chronic diseases such as emphysema/chronic obstructive pulmonary disease (COPD), idiopathic pulmonary fibrosis (IPF), and bronchopulmonary dysplasia (BPD), characterized by impaired or arrested alveolar development, as well as the acute respiratory distress syndromes accompanying severe injury or alveolar damage following infection, as in SARS, MERS, and the current Covid-19 pandemic (*19–24*). In these diseases enlargement, destruction, or flooding of alveoli, or changes in thickness or composition of their walls, result in a loss or lack of gas exchange surface, causing a decline in lung function or even respiratory failure.

The textbook model of alveolar development posits that the epithelium forms large sacs (‘sacculation’) that are later subdivided into alveoli by contractile myofibroblasts (‘septation’) (Fig. S1) (*25–29*). Alveolar septa are thought to arise postnatally in rodents (*25, 27*) and beginning at fetal stages in humans (*30, 31*), initially as shallow crests that then deepen, with myofibroblasts (first termed ‘contractile interstitial cells’ because of their characteristic ultrastructure and contractile elements similar to smooth muscle) and elastic fibers located at septal tips (*32–35*). Evidence to support the model is limited since developing alveoli have not been visualized at cellular resolution and in three dimensions, and the origin and role of myofibroblasts are not well defined. Although recent studies have begun to localize and characterize myofibroblasts and their progenitors (*36–39*), it remains unclear when and how myofibroblasts arise and are recruited to epithelial sacs, how they get positioned or anchored on the saccular walls, and how they migrate inward or contract to produce alveoli. Despite remarkable recent progress in alveolar development (*40–45*) including dynamic cell imaging (*46*) and elucidation of the complete gene expression program of the alveolar epithelium (*47*), our understanding of the mechanisms of alveolar patterning and morphogenesis and the coordination and interactions between cells remains rudimentary.

Here we use mosaic labeling, clonal analysis and deep imaging to elucidate the three-dimensional cellular structure of alveoli in the mouse lung. We then analyze alveolar development at single-cell resolution and at a specific position in the branch lineage. We show that alveoli form much earlier and more simply than previously thought – by budding of the epithelium on airway stalks between smooth muscle cells that rearrange to become myofibroblasts at the alveolar openings. A single AT2 cell is selected to lead the alveolar outgrowth; a lateral inhibitory signal mediated by Notch prevents induction of AT2 fate in surrounding progenitors, which then develop as AT1 cells. Our analysis suggests an elegant new mechanism of alveolar patterning and development distinct from the classical septation model.

### The three-dimensional cellular structure of alveoli in the mouse lung

To visualize the three-dimensional architecture of individual alveoli, we labeled the network of elastic fibers that surround and support the entrance to alveoli (*48, 49*) by incubating inflated and fixed lungs in fluorescent hydrazide (Fig. S2) (*50*). Deep imaging and three-dimensional reconstruction revealed three classes of alveolar elastic fibers delineated by thickness and localization within or around alveoli (thick, thin and microvascular fibers; Fig. S2C). These classes allowed us to discern two types of alveoli: simple alveoli with the entrance formed by a ring of thick fibers (Fig. 1, A and B, Fig. S2D, and Movie S1), and groups of 2-8 alveoli (alveolar complexes) with a shared entrance encircled by thick fibers and partitions formed by thin fibers (Fig. 1, C and D, Fig. S2E, and Movie S2). This is consistent with previous descriptions of the geometry of respiratory airways (*1, 2, 8, 51*). Single alveolar units and complexes are intermingled throughout respiratory airways, although simple outpocketings are especially obvious and may be more abundant on early generations (Fig. S2, F and G).

**Figure 1.**
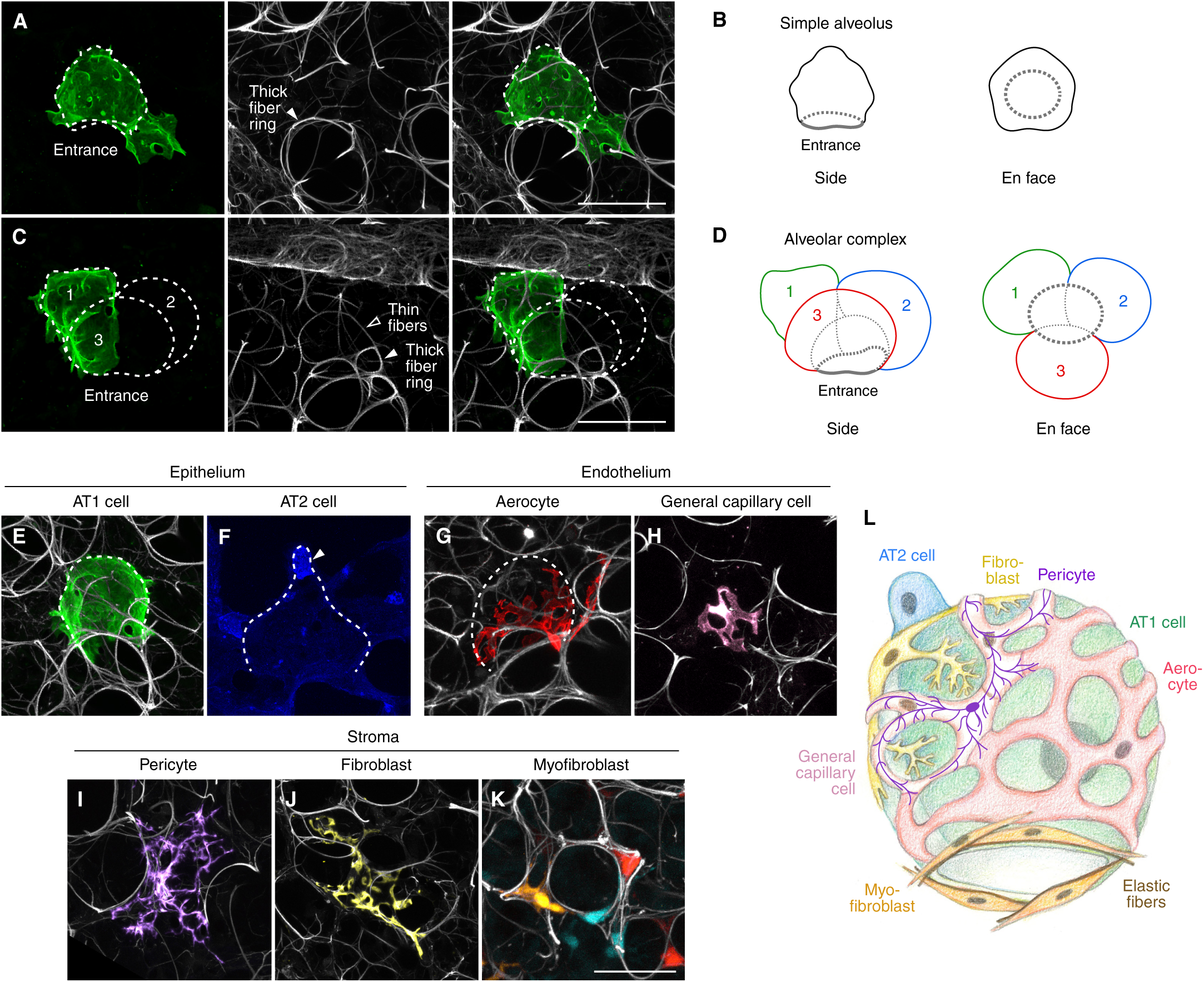
Single-cell labeling reveals the cellular structure of alveoli in the mouse lung. (**A**, **B**) Simple alveolus. Confocal projection (**A**) and diagram (**B**) of a simple alveolus (dashed outline) with genetic labeling of single AT1 cell (green) in a *Hopx-CreER; Rosa26-mTmG* lung dosed with limiting tamoxifen (0.3 mg) 5 days prior to analysis at 2 months of age and co-stained with GFP antibodies to visualize labeled cells and fluorescent hydrazide to label elastic fibers (white). Thick elastic fibers (solid arrowheads in **A**) form a ring at the alveolar entrance (dotted grey circles in **B**). Diagram also shows en face view (right) of same alveolus. See also Movie S1. Scale bars, 50 µm. (**C**, **D**) Alveolar complex. Confocal projection (**C**) and diagram (**D**) as above of an alveolar complex. Thin elastic fibers (open arrowheads in **C**, dotted grey lines in **D**) subdivide the alveolar complex into 3 units, which are numbered and outlined in green, blue, and red in **D**. See also Movie S2. Scale bars, 50 µm. (**E** to **K**) Confocal projections showing morphologies and arrangement of single alveolar cells [AT1 cell (**E**, green), AT2 cell (**F**, blue, arrowhead), capillary aerocyte (**G**, red), general capillary cell (**H**, pink), pericyte (**I**, purple), fibroblast (**J**, yellow), and myofibroblasts (**K**, labeled with the multicolor fluorescent reporter *Rosa26-Rainbow*)] in alveoli (dotted outlines in **E** to **G**) and relative to elastic fibers (white, **E**, **G** to **K**) in a *Hopx-CreER; Rosa26-mTmG* lung (**E**) dosed and analyzed as above, a *Shh-Cre; Rosa26-mTmG* lung (**F**) immunostained for GFP at 2 months, a *VE-cadherin-CreER; Rosa26-Confetti* lung (**G** and **H**) dosed with 1 mg tamoxifen at 3 weeks and analyzed at 2 months by imaging endogenous RFP fluorescence, a *Pdgfra-CreER; Rosa26-mTmG* lung (**I**) dosed with 2 mg tamoxifen at 3 months and immunostained for GFP, a *Gli1-CreER; Rosa26-mTmG* lung (**J**) dosed with 4 mg tamoxifen at 2 months and immunostained for GFP, and a *SMMHC-CreER; Rosa26-Rainbow* (**K**) lung dosed with 0.1 mg tamoxifen at P5, immunostained at P10 with SMA-Cy5 antibodies (not shown) and analyzed by imaging endogenous Cerulean, mCherry and mOrange fluorescence. Scale bar, 50 µm. (**L**) Schematic of an alveolus with 7 major cell types (not showing immune cells) and a total of 12 cells: a single AT1 cell (green), an AT2 cell (blue), 3 capillary cells (1 aerocyte, red; 2 general capillary cells, pink; (*53*)), 2 pericytes (purple, with 1 on backside not shown), 1 fibroblast (yellow), and 4 myofibroblasts (orange) that associate with elastic fibers (brown) at the alveolar base. Grey ovals, nuclei.

To elucidate the cellular structure of alveoli, we determined the morphologies of alveolar cells and their arrangement by labeling single cells using genetic strategies and analyzing their position within alveoli relative to the ring of elastic fibers at the alveolar entrance. Tamoxifen-inducible CreER lines (Table S1), in which Cre recombinase is fused to a modified estrogen receptor to control the timing and extent of labeling, were crossed to a Cre reporter in which recombination of a *loxP* flanked stop cassette leads to expression of a fluorescent protein. Limiting doses of tamoxifen were used to induce rare recombination events (100-800 events per lobe) that labeled isolated cells. Three-dimensional reconstruction of single cells and the surrounding fibers revealed that alveolar cells have a stunning variety of shapes and unusual features and arrangements (Fig. 1, E to K).

There is on average just a single alveolar type 1 (AT1) cell per alveolus (Fig. 1E, Fig. S3, and Fig. S5, A, B and G). It is typically a cup-shaped cell with its thin, expansive cytoplasmic extensions covering nearly the entire alveolar surface (5,500 µm^2^) (*12*). However, the arrangement of AT1 cells in alveoli is variable and individual cells can span 2-5 alveoli (Fig. S3 and S4), consistent with descriptions of ‘non-nucleated plates’ (*17*). The majority (96%, n=100) of AT1 cells have one or more (up to 9) holes (‘pores of Kohn’ (*52*)) of circular or oval shape and varying sizes (Fig. S5B). AT1 cells form junctions with each other and with neighboring AT2 cells that are strategically positioned between alveoli as single cuboidal or elongated cells (Fig. 1F and Fig. S5C) (*17*). Although there is typically also just a single AT2 cell per alveolus (Fig. S5G), individual AT2 cells often have multiple apical surfaces contributing to 2-4 alveoli (80% of AT2 cells with >1 apical domain, n=500).

The alveolus is surrounded by a dense mesh of capillaries made up of two distinct, intermingled cell types with ‘swiss cheese’ morphologies (Fig. 1G and H, and Fig. S6, A and B) (*53, 54*). Pericytes and alveolar fibroblasts are complex, branched cells with long (>50 µm), thin processes spanning multiple alveoli (Fig. 1, I and J, Fig. S5, E and F, and Fig. S6, C and D). Myofibroblasts, by contrast, are simple, spindle-shaped cells with 2 or 3 cell processes that are aligned with thick elastic fibers at the alveolar entrance (Fig. 1K and Fig. S5D). Three to five myofibroblasts form a ring around the alveolar entrance (Fig. S5G), with their cell bodies located at the intersections of neighboring alveoli and with the number of cells per ring corresponding to the number of intersections.

These clonal labeling experiments combined with immunostaining against common antigens, deep imaging, 3D reconstruction and quantification (Figs. S3-S6 and Methods) allowed us to define the three-dimensional cellular structure of an alveolus (Fig. 1L and Fig. S5G). A simple alveolus is composed of just 2 epithelial cells, a single AT2 cell and an AT1 cell that covers the majority of the alveolar surface, 3-4 capillary endothelial cells (of 2 cell types: aerocytes and general capillary cells; relative abundance 1:4), and 5-9 stromal cells including 1-2 pericytes, 1-2 fibroblasts, and 3-5 myofibroblasts. Thus, an alveolus in the mouse lung is composed of just 10-15 cells of 7 major types, excluding immune cells. This is 2-3 fold fewer cells than previous estimates based on stereology of thin sections of alveoli (*55*), which apparently did not distinguish complexes from individual alveoli. Alveolar complexes contain proportionally more cells than simple alveoli, scaled to the number of alveolar units (“chambers”) in the complex (Fig. S4).

### Mapping alveolar development at cellular resolution at a defined position in the lung

To elucidate the cellular mechanisms that initiate and control alveolar patterning, we analyzed the earliest events in alveolar development by mapping the process at a specific position in the branch lineage (*56*). In the left primary bronchus (L) lineage, the third anterior branch (A3) forms by domain branching on the anterior aspect of the first secondary branch (L1) and generates 4-6 generations of daughter branches by planar and orthogonal bifurcations (Fig. 2A). By immunostaining of whole lobes for Sox2 (or incubation with fluorescent streptavidin to detect biotin expressed by epithelial cells), which allows visualization of conducting airways (*57, 58*), we mapped the boundary between conducting and respiratory airways on L.L1.A3 and analyzed when the boundary forms during development (Fig. S7). We found that the boundary is positioned on L.L1.A3 daughters at or just distal to the bifurcation between airway generations 4 and 5 (Fig. S7, A to C). Consistent with previous studies (*57*), the boundary is set by embryonic day (E) 16.5 (Fig. 2, B to D, and Fig. S7D), and branches distal to the boundary form respiratory airways.

**Figure 2.**
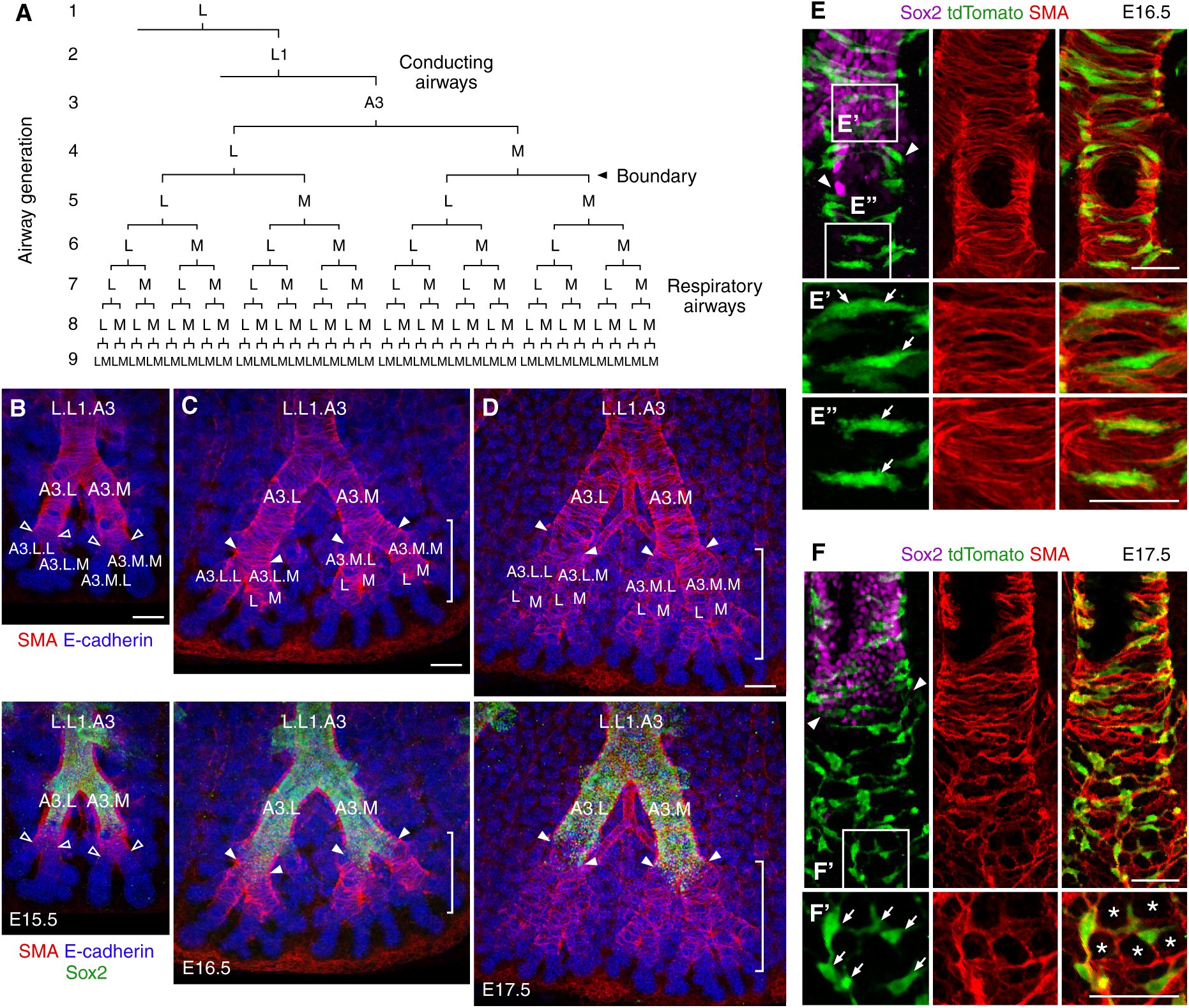
Mapping alveolar development at a specific position in branch lineage. (**A**) Branch lineage diagram for third anterior branch of left primary bronchus (L.L1.A3) showing descendant branches and position of the boundary (arrowhead) between conducting and respiratory airways along branches of bronchial generations 4-5. Airway generations are numbered. A, anterior; L, lateral; M; medial. (**B** to **D**) Confocal projections showing L.L1.A3 branch lineage in wild-type lungs immunostained for E-cadherin (epithelium, blue), SMA (smooth muscle, red) and Sox2 (conducting airways, green) at the indicated embryonic ages. The boundary between conducting and respiratory airways (solid arrowheads in **C**, **D**) is set on L.L1.A3 daughters by E16.5. Branches distal to the domain marked by Sox2 (brackets in **C**, **D)** form respiratory airways; their names are indicated in top panels. SMA-positive cells (red) cover airway stalks and a few cells extend beyond the inferred future boundary position (open arrowheads in **B**) at E15.5. Once the boundary is set, stalks of respiratory airways are surrounded by a layer of SMA-positive cells (brackets in **C**). Note that the orientation of cells on airway stalks changes one day later (brackets in **D**). Scale bars, 50 µm. (**E**) Confocal projection with close-ups of boxed regions (**E’** to **E’’**) showing SMA-positive cells (arrows in **E’** to **E’’**) on stalks of conducting (**E’**) and respiratory airways (**E’’**) in an E16.5 *SMA-CreER; Rosa26-tdTomato* lung dosed with tamoxifen (2 mg) at E15.5 and immunostained for Sox2 (conducting airways, magenta), SMA (smooth muscle, red) and the lineage tag (tdTomato, pseudocolored in green). Note similar morphology and circumferential orientation of the cells at both positions at this stage. Arrowheads, compartment boundary. Scale bars, 50 µm. (**F**) Confocal projection with close-up of boxed region (**F’**) as above in panel **E** showing SMA-positive cells (arrows in **F’**) on stalks of respiratory airways in an E17.5 *SMA-CreER; Rosa26-tdTomato* lung dosed with tamoxifen (2 mg) at E15.5. The lineage tag (tdTomato) is pseudocolored in green. Note that at this stage (E17.5) the cells have reoriented and form rings (asterisks in **F’**) on stalks of respiratory airways. Scale bars, 50 µm.

Immunostaining of whole lobes for epithelial and smooth muscle markers showed that at E16.5 smooth muscle actin (SMA)-positive cells surround epithelial tubes destined to form respiratory airways (Fig. 2C). The cells have an elongated morphology and circumferential orientation typical of smooth muscle on conducting airways (Fig. 2E). Just one day later (E17.5), the smooth muscle pattern is less organized (Fig. 2D), with the cells oriented along airway stalks at various angles (Fig. 2F). Already at this stage, the cells are aligned with elastic fibers (Fig. S8). This suggested that smooth muscle cells on embryonic airways rearrange and become myofibroblasts at the alveolar entrance.

### Airway smooth muscle cells become myofibroblasts at the alveolar entrance

To test whether airway smooth muscle is a source of myofibroblasts, we used a genetic strategy to specifically label and trace smooth muscle on embryonic airways (*59*). We combined the airway smooth muscle-specific *Lgr6-EGFP-IRES-CreERT2 (Lgr6-CreER)* (*60*) knock-in allele with the Cre reporter *Rosa26-tdTomato* (*61*) and induced recombination at E14.5 with a saturating dose of tamoxifen (Fig. 3A and Fig. S9). Analysis of the labeling on postnatal day 10 (P10) showed lineage-labeled cells on first and second order respiratory airways (Fig. 3B). The cells express SMA and are aligned with thick elastic fibers at alveolar openings (Fig. 3B’), demonstrating that airway smooth muscle cells become entrance myofibroblasts. Alveolar entrance myofibroblasts express *Eln* (tropoelastin) as shown by single-cell RNA-sequencing (scRNAseq) and single-molecule in situ hybridization (smFISH) (Fig. S10, F and G) (*62, 63*), confirming that the cells are indeed a source of elastin, a major component of elastic fibers.

**Figure 3.**
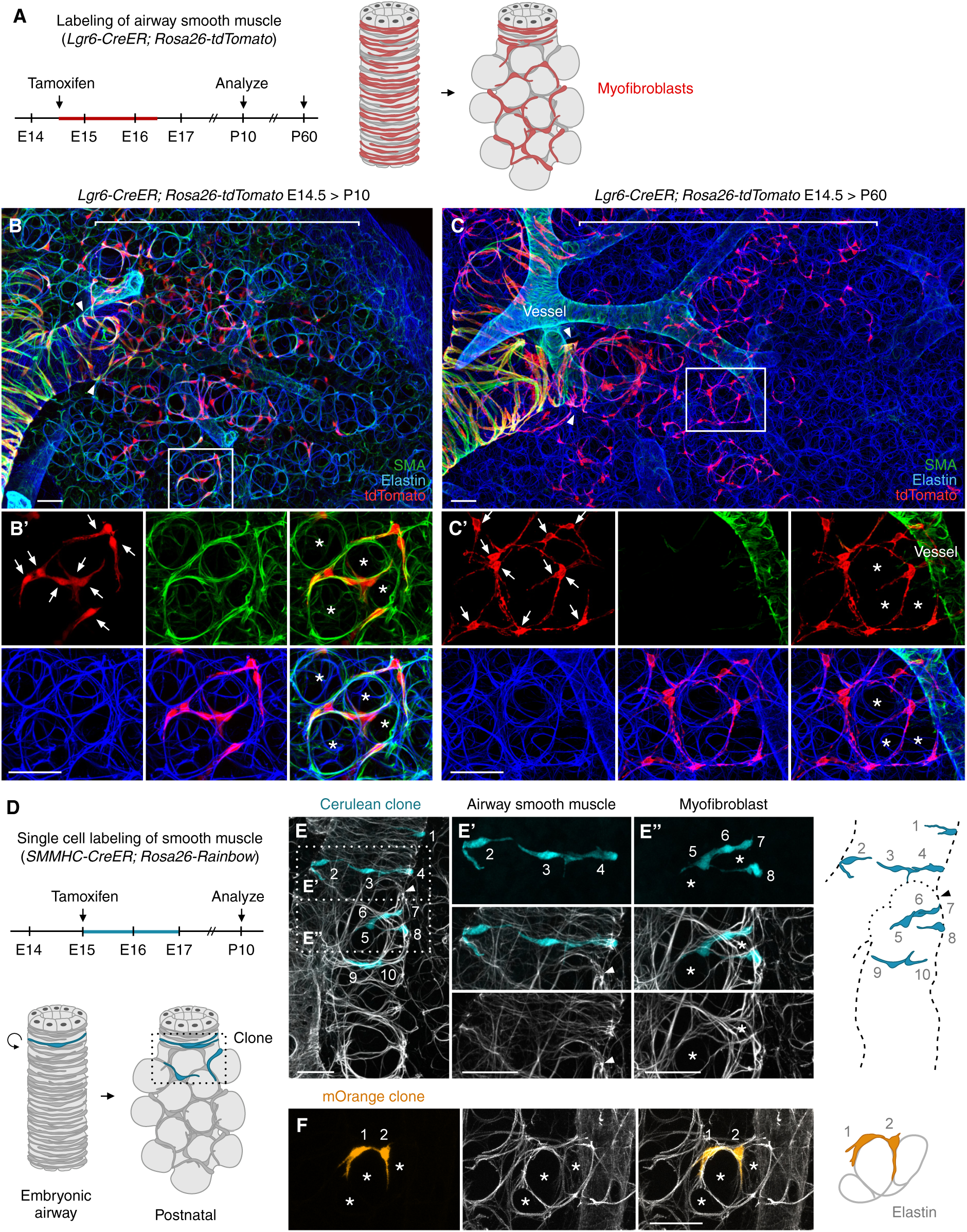
Lineage tracing and clonal analysis show that airway smooth muscle cells become alveolar myofibroblasts. (**A**) Strategy of lineage trace of airway smooth muscle. *Lgr6-CreER; Rosa26-tdTomato* lungs were dosed at E14.5 with saturating (4 mg) tamoxifen to label smooth muscle on airway stalks with heritable tdTomato expression (red bar on diagram, with tamoxifen assumed active for 48 hours). Labeling (left panel of schematic) was analyzed at P10 and P60 once alveoli have formed (right panel of schematic). (**B** and **C**) Confocal projections with close-ups of boxed regions (**B’**, **C’**) showing alignment of lineage-labeled cells (arrows in **B’**, **C’**) with elastic fibers (blue) at the alveolar entrance in *Lgr6-CreER; Rosa26-tdTomato* lungs dosed as above and stained at P10 (**B**) or P60 (**C**) for the lineage tag (tdTomato, red), SMA (smooth muscle actin, green) and elastin (detected by hydrazide; blue). Asterisks in **B’** and **C’** denote alveolar lumens. Lineage-labeled alveolar cells express SMA at P10 (**B’**), but not at P60 (**C’**). Arrowheads, compartment boundary (identified by staining with fluorescent streptavidin to detect epithelial biotin; not shown). Brackets, respiratory airways with lineage-labeled cells. Lineage-labeled cells are located on 2 generations of respiratory airways, which corresponds to the number of airway generations surrounded by smooth muscle beyond the compartment boundary at E16.5 (see Fig. 2C). Incomplete labeling of alveolar entrance rings on these airways may be due to inefficient recombination in airway smooth muscle (see Fig. S9). Scale bars, 50 µm. (**D**) Strategy of single cell (clonal) labeling of smooth muscle. *SMMHC-CreER; Rosa26-Rainbow* lungs were dosed at E15 with limiting doses (0.05 mg) of tamoxifen to label single smooth muscle cells (bottom diagram, left) in one of 3 colors (Cerulean, mCherry, mOrange). Labeling was analyzed around airways at P10 (bottom diagram, right). Blue bar, labeling window (tamoxifen assumed active for 48 hours). Vascular smooth muscle clones were not analyzed. (**E**) Confocal projection (left) with close-ups of boxed regions (**E’**, **E’’**) and diagram (right) of a mixed clone (Cerulean) generated with *SMMHC-CreER; Rosa26-Rainbow* as described in (**D**) with labeled airway smooth muscle cells (**E’**) and myofibroblasts (**E’’**) that are aligned with rings of elastic fibers (white) at the alveolar entrance. Asterisks denote alveolar lumens. Cells in the clone are numbered. Arrowheads, compartment boundary. Scale bars, 50 µm. (**F**) Confocal projection (left) and diagram (right) of myofibroblast clone (mOrange) generated with *SMMHC-CreER; Rosa26-Rainbow* as above. Cells in the clone are numbered. Two myofibroblasts (orange) are aligned with elastic fiber rings (white) at alveolar openings. Note incomplete labeling of entrance rings. Scale bars, 50 µm.

To determine if myofibroblasts on later-generation branches also arise from the Lgr6 lineage, we induced recombination at a later embryonic timepoint by dosing E17.5 *Lgr6-CreER; Rosa26-tdTomato* lungs with tamoxifen (Fig. S10A). Lineage labeled cells were located on 3-4 generations of respiratory airways (Fig. S10, B and B’). Postnatal labeling (Fig. S10C) included distal respiratory airways (Fig. S10, D and D’). This shows that alveolar entrance myofibroblasts are derived from the Lgr6 lineage (Fig. S10H) on early and late generation airways.

To probe the potential and behavior of individual airway smooth muscle cells, we sparsely labeled smooth muscle cells by induction of rare recombination events in smooth muscle myosin heavy chain *(SMMHC)-CreER; Rosa26-Rainbow* lungs (*64, 65*) using limiting doses of tamoxifen (Fig. 3D). Airway clones induced at E15 and analyzed on postnatal day 10 were composed of 1 to 17 cells. Clones fell into 3 groups based on the location of daughter cells (Table S2). In the first group, cells of a clone were located exclusively around stalks of conducting airways (“airway smooth muscle clones”, n=30 clones; 44%; Fig. S11). In some of these clones (37%) the daughter cells remained in close contact with their siblings, whereas in other clones (37%) cells were dispersed, spanning 100-500 µm along the airway axis, implying extensive movement of daughter cells, as has been observed early in development (*59*). The second group was formed by clones spanning the boundary between conducting and respiratory airways (“mixed/boundary clones”, n=8 clones; 12%; Fig. 3E and Fig. S12). These clones were composed of both elongated cells with circumferential orientation around airway stalks (airway smooth muscle) and spindle-shaped cells aligned with elastic fibers at the alveolar entrance (myofibroblasts). This demonstrates that individual cells can give rise to both, establishing a lineage relationship between airway smooth muscle cells and myofibroblasts. The third group of clones was composed of cells located exclusively on respiratory airways (“myofibroblast clones”, n=30 clones; 44%; Fig. 3F and Fig. S13). Nearly all (98%) labeled cells (n=100 cells scored from 30 clones) were aligned with alveolar elastic fiber rings. Although the majority (70%) of myofibroblast clones were composed of more than one cell (Table S2), individual entrance rings were generally incompletely labeled, excluding a model of entrance ring formation by clonal proliferation. The differences in cell behavior our analysis uncovered, with only a fraction of labeled cells turning into myofibroblasts, may reflect differences in the potential of individual cells, or perhaps more likely be dictated by the environment (e.g., only daughter cells located beyond the boundary turn into myofibroblasts).

### Anatomical and developmental diversity of alveolar myofibroblasts

Immunostaining and smFISH in postnatal (P10) lungs revealed two distinct types of myofibroblasts. Both types express SMA and the myofibroblast marker *Fgf18* (*66*), but one population associates with thick elastic fibers that encircle the openings to simple alveoli and alveolar complexes (“entrance myofibroblasts”; Fig. 4, A, A’, B, B’ and C). Our embryonic and postnatal pulse-chase experiments with *Lgr6-CreER; Rosa26-tdTomato* showed that entrance myofibroblasts derive from the Lgr6 lineage (Fig. 3B’ and Fig. S10, A to D) and persist in the adult lung (Fig. 3C and Fig. S10E). Although nearly all (98%) of these cells turn off SMA, they remain aligned with thick elastic fibers and retain a simple morphology (Fig. 3C’).

**Figure 4.**
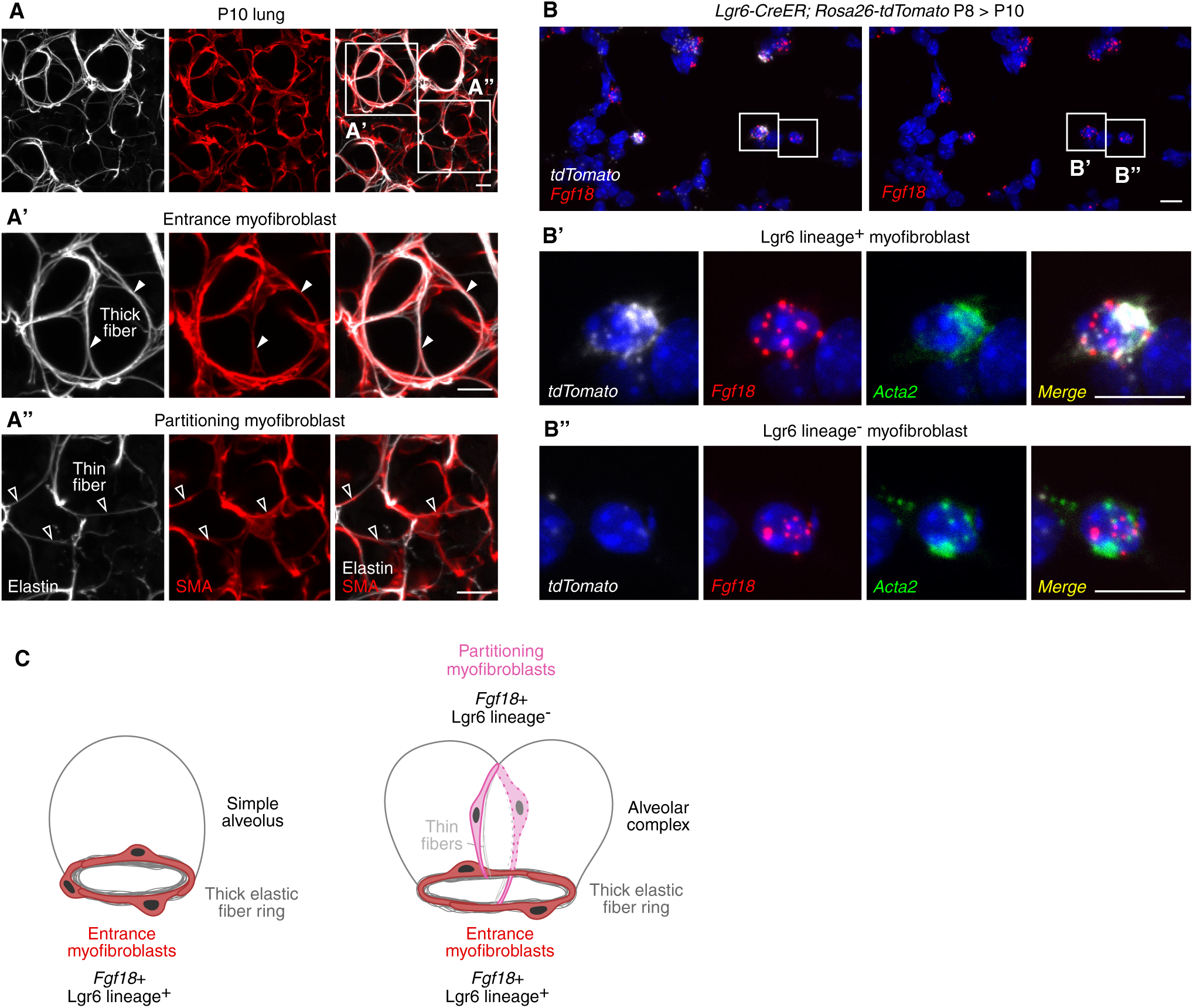
Anatomical and developmental diversity of myofibroblasts. (**A**) Confocal projections with close-ups of boxed regions (**A’** and **A’’**) of a P10 C57BL/6 lung stained for SMA (smooth muscle actin, red) and elastin (detected by hydrazide, white). Filled arrowheads, alveolar entrance myofibroblasts aligned with thick elastic fibers; open arrowheads, partitioning myofibroblasts aligned with thin fibers. Scale bars, 10 µm. (**B**) Single-molecule in situ hybridization for *tdTomato* (white), *Fgf18* (red) and *Acta2* (SMA, green) in P10 *Lgr6-CreER; Rosa26-tdTomato* lung labeled at P8. *Fgf18*+ *tdTomato*+ cells (Lgr6 lineage-derived entrance myofibroblasts) (**B’**) comprise a subset of *Acta2*+ cells (21 ± 4%; mean ± s.d.; n=3 lungs). The majority (98 ± 1%) of entrance myofibroblasts (Lgr6 lineage-labeled cells) express *Fgf18* and *Acta2*. Partitioning myofibroblasts (**B’’**) also express *Fgf18* and *Acta2*. Scale bars, 10 µm. (**C**) Schematic of entrance myofibroblasts (red) in simple alveoli (left) and alveolar complex (right) and partitioning myofibroblasts (pink) that subdivide alveolar complex into two units.

The second population also expresses SMA and the myofibroblast marker *Fgf18*, but these cells are aligned with thin elastic fibers that subdivide alveolar complexes (“partitioning myofibroblasts”, Fig. 4, A, A’’, B, B’’ and C). Partitioning myofibroblasts also have a spindle shape, and their cell processes are anchored on the thick elastic fibers and entrance myofibroblasts at the openings to alveolar complexes. Unlike alveolar entrance myofibroblasts, partitioning myofibroblasts are not lineage labeled by *Lgr6-CreER* (Fig. 3B’ and Fig. S10, A to D), hence they arise from another source.

These results suggest that there are two types of myofibroblasts with distinct anatomical locations and developmental origins: SMA+ *Fgf18*+ Lgr6 lineage-derived entrance myofibroblasts that form the thick fiber rings at the base of simple alveoli and alveolar complexes, and SMA+ *Fgf18*+ Lgr6 lineage-negative partitioning myofibroblasts that subdivide alveolar complexes. Simple alveoli have entrance but not partitioning myofibroblasts (Fig. 4C).

### Alveoli form by epithelial budding led by a single AT2 cell

The alignment of SMA+ cells with thick elastic fiber rings on stalks of embryonic airways at E17.5 (Fig. S8) suggested that alveoli form much earlier than previously assumed (P4 in mice) (*25*) and by a mechanism distinct from septation.

To determine how alveoli arise on airways, we analyzed the epithelial pattern on stalks of L.L1.A3 daughter branches by staining embryonic (E17.5) lobes for E-cadherin, and also for SMA and elastin (detected by hydrazide) to visualize the arrangement of epithelial cells relative to smooth muscle and elastic fibers. Surprisingly, we found small groups of 1-3 cuboidal epithelial cells protruding between smooth muscle cells on airway stalks (Fig. 5A and Fig. S14). The budding single cuboidal cells were surrounded by rings of 3 or more smooth muscle cells aligned with elastic fibers (Fig. 5, A’ and A’’), indicating that these are the nascent entrance myofibroblasts organizing around budding epithelial cells and depositing elastin, supporting a model of alveolar development by direct budding.

**Figure 5.**
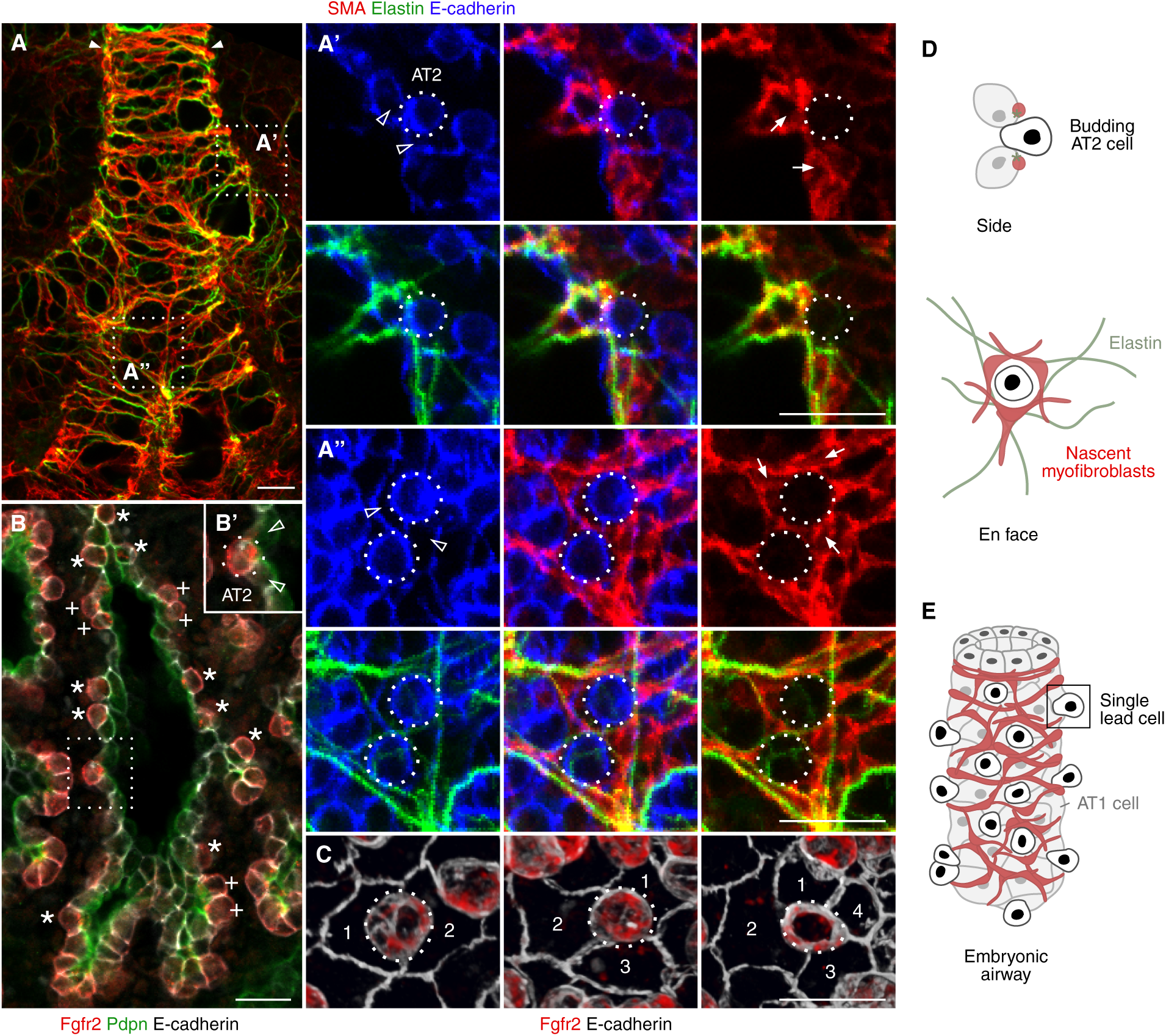
Alveolar buds are led by single AT2 cells through nascent myofibroblasts. (**A**) Confocal projection with close-ups of boxed regions (**A’**, side view and **A’’**, en face view) showing that single cuboidal epithelial cells (dotted circles) are surrounded by partially flattened cells (open arrowheads) and a ring formed by smooth muscle cells (arrows) that are aligned with elastic fibers (green) in an E17.5 wild-type lung stained for SMA (smooth muscle actin, red), elastin (detected by hydrazide, green) and E-cadherin (epithelium, blue). Filled arrowheads, compartment boundary. Scale bars, 20 µm. (**B**) Confocal slice with close-up of boxed region (**B’**) of an E17.5 wild-type lung immunostained for FGF receptor 2 (Fgfr2, progenitors and AT2 cells, red), podoplanin (Pdpn, progenitors and AT1 cells, green), and E-cadherin (epithelium, white) showing that alveolar epithelial buds are led by single AT2 cells (asterisks; note Fgfr2 restriction to single cuboidal epithelial cells, or occasionally, groups of 2-3 cuboidal cells, denoted by crosses). Single AT2 cells (dotted circle in **B’**) are surrounded by nascent AT1 cells (open arrowheads in **B’**). Clustered cuboidal cells in distal tips are progenitors (Fgfr2+ Pdpn+ cells) that have not yet undergone differentiation into AT2 or AT1 cells. (**C**) Confocal projections of an E18.5 wild-type lung immunostained for Fgfr2 (red) and E-cadherin (epithelium, white) showing regular salt-and-pepper pattern of the alveolar epithelium. Nascent AT2 cells are surrounded by 2-4 partially flattened cells that do not express Fgfr2 (nascent AT1 cells, numbered). Scale bars, 20 µm. (**D**) Diagrams of nascent alveolar buds, shown as side (top) and en face views (bottom), led by a single AT2 cell (black) through the nascent myofibroblasts (red) that synthesize and align with elastic fibers (green). (**E**) Diagram of airway with leading single AT2 cells (black) surrounded by AT1 cells (light grey) and ring of myofibroblasts (red).

Immunostaining for alveolar epithelial markers, including FGF receptor 2 (Fgfr2), showed that the single cuboidal cells are nascent AT2 cells (Fig. 5B and Fig. S15; see also Brownfield et al., in review (*67*)). Each budding AT2 cell is surrounded by 2-4 nascent AT1 cells that have begun to flatten (Fig. 5C). This suggests that the alveolar buds are led by single AT2 cells through the nascent entrance myofibroblasts that begin to synthesize elastic fibers and demarcate the alveolar entrance as the AT2 cell buds (Fig. 5D and E).

### A lateral inhibitory signal mediated by Notch ensures selection of a single AT2 cell

The regular salt-and-pepper pattern of the alveolar epithelium with single AT2 cells surrounded by 2-4 nascent AT1 cells implies that the alveolar pre-pattern and spacing is established surprisingly early in the process, at or before E17.5. The salt-and-pepper pattern also raised the possibility that the balance between the alternative epithelial fates could be controlled by a Notch-mediated lateral inhibition mechanism (*68*). Indeed, an increase in AT2 cells has been observed in embryonic lungs mutant for *Lunatic Fringe*, a glycosyltransferase that facilitates Notch activation (*69*), and Notch signaling is active during the AT2-to-AT1 cell transition in culture (*70*). To determine if the Notch pathway is active in developing alveolar epithelial cells, we examined expression of the major transcriptional effector of Notch signaling, CBF1 (RBP-J), in embryonic (E16.5-18.5) lungs using a *CBF1-H2B-Venus* reporter (*71*). Immunostaining showed nuclear Venus localization in nascent AT1 cells (and endothelial plexus) but little or none in alveolar epithelial progenitors or budding AT2 cells marked by Fgfr2 (Fig. 6A and Fig. S16, A and B). Immunostaining for the Notch target Hes1 also showed pathway induction in nascent AT1 cells (Fig. S16, C and D). Thus, the Notch pathway is activated in the AT1 lineage during alveolar development.

**Figure 6.**
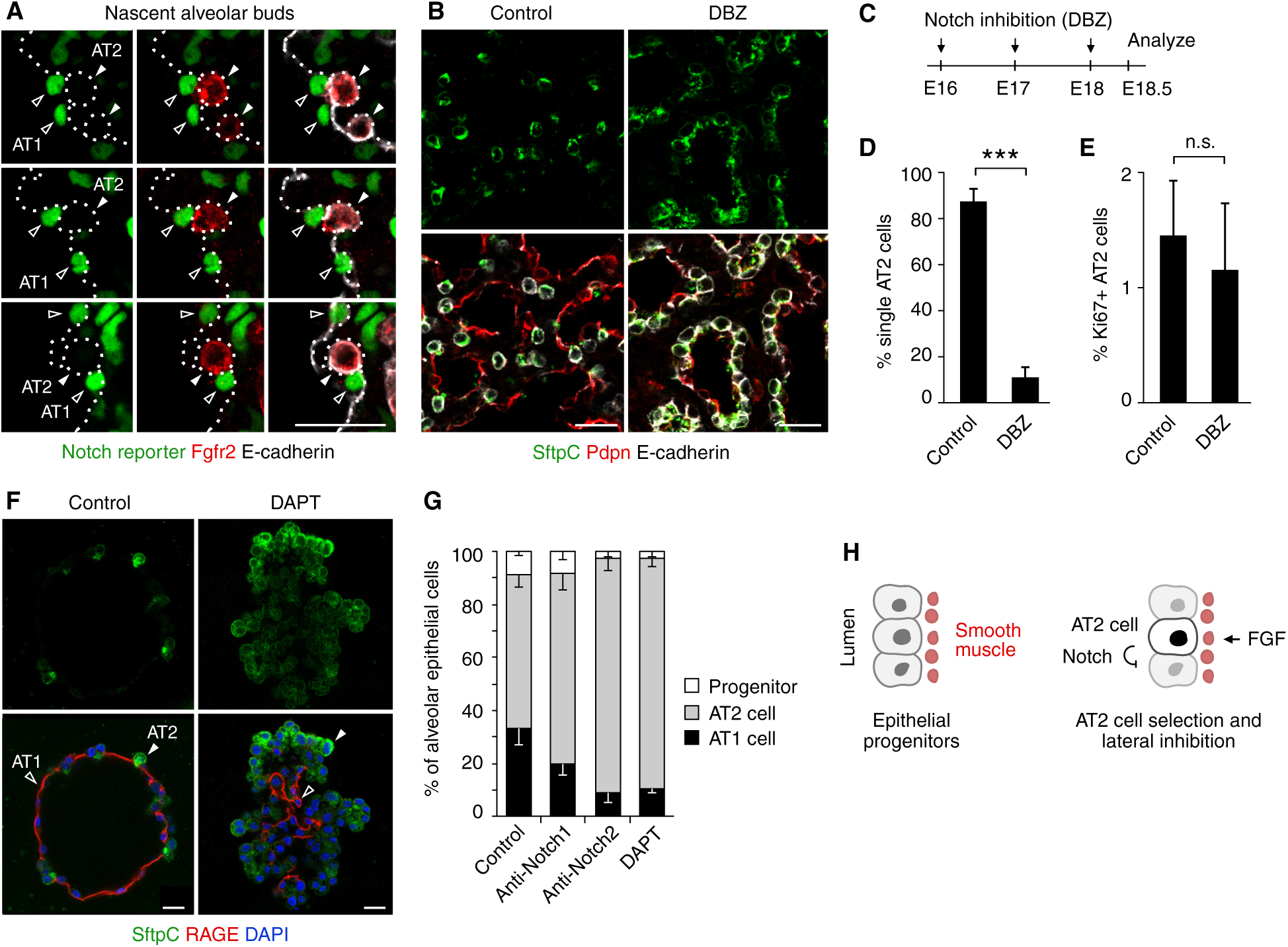
Effect of Notch signaling inhibition on alveolar epithelial patterning. (**A**) Single confocal slices showing nascent alveolar buds of an E17.5 *CBF1-H2B-Venus* lung immunostained for GFP to detect Venus (Notch nuclear reporter, green), Fgfr2 (red), and E-cadherin (epithelium, white). Nascent AT2 cells (dotted circles and solid arrowheads) express Fgfr2, but not Venus. Neighboring AT1 cells (dotted lines and open arrowheads) express nuclear Venus, but not Fgfr2. (**B**) Effect of pharmacological Notch inhibition by DBZ on alveolar epithelial patterning. DBZ (30 µM per kilogram body weight) or vehicle were injected daily in wild-type mice (E16-E18, panel **C**). Single confocal slices of vehicle (control) or DBZ-treated E18.5 lungs immunostained for SftpC (progenitors and AT2 cells, green), podoplanin (Pdpn, progenitors and AT1 cells, red) and E-cadherin (epithelium, white). (**C**) DBZ dosing regime. (**D**) Quantification of AT2 cell clustering in E18.5 vehicle or DBZ-treated lungs (mean ± s.d.; n=500 cells scored per mouse; 6 mice per treatment group; Wilcoxon rank sum test, p-value=0.002 (***)). (**E**) Quantification of AT2 cell proliferation in E18.5 vehicle or DBZ-treated lungs (mean ± s.d.; n=500 cells scored per mouse; 3 mice per treatment group; Wilcoxon rank sum test, p-value=0.69 (n.s., not significant) comparing % Ki67+ AT2 cells). (**F**) Effect of Notch inhibition on fate selection of alveolar epithelial progenitors isolated from E16.5 wild-type lungs and cultured in Matrigel with Fgf7 and Notch inhibitor DAPT or Notch1 or Notch2 blocking antibodies as indicated. Single confocal slices of cultures on day 4 immunostained for SftpC (progenitors and AT2 cells, green) and RAGE (progenitors and AT1 cells, red) and counterstained with DAPI (nuclei, blue). (**G**) Quantification of alveolar cell fate (mean ± s.d.; n=500 cells scored in 3 biological replicates). Scale bars, 20 µm. (**H**) Schematic of Notch-mediated lateral inhibition of AT2 fate following AT2 induction by Fgf signaling.

To test if Notch signaling plays a role in alveolar patterning, we inhibited the pathway in developing (E16-E18.5) lungs using the γ-secretase inhibitor DBZ, which prevents ligand-induced proteolytic processing and activation of Notch (*72*). Whereas the majority (88 ± 5%) of AT2 cells in control lungs (n=500 AT2 cells scored in 6 animals) were found as single (isolated) AT2 cells at E18.5, only rare AT2 cells (11 ± 4%) were selected as single cells in DBZ-treated lungs (n= 500 cells in 6 animals), with the rest (89 ± 4%) found in clusters (Fig. 6, B to D). To determine if Notch inhibition enhances AT2 cell proliferation, as observed at postnatal stages (*73*), we performed immunostaining for Ki67. AT2 cell proliferation was comparable in DBZ-treated and control lungs (1.2 ± 0.6% Ki67+ AT2 cells in DBZ-treated lungs; 1.5 ± 0.5% in controls; Fig. 6E and Fig. S16E). This suggests that Notch signaling is required for alveolar patterning and specifically for selection of single AT2 cells, rather than to regulate their proliferation. Technical, developmental timing or background differences could explain the discrepancy between our findings and a previous study of Notch function (*73*).

To identify the relevant receptor and ligand, we analyzed expression of Notch pathway components in scRNAseq data for developing mouse lung (*47, 74*). This confirmed Notch pathway activation (assessed by *Hes1* expression) in the AT1 lineage. However, it also revealed potential signaling complexity. Expression of multiple receptors (*Notch1-4*) and ligands (*Jag1, Jag2, Dll1, Dll4*) was detected in the developing alveolar epithelium (Fig. S17, A and B) along with non-canonical ligands *Dlk1* and *Dlk2*, with *Dlk1* induced in the AT2 lineage. Other (non-epithelial) alveolar cells also expressed ligand genes (Fig. S17, B and C) identifying other possible sources of signal.

To test for Notch-mediated lateral inhibition, we used a cell culture system in which alveolar epithelial progenitors from E16.5 lungs are purified away from other alveolar cells and cultured alone in Matrigel; addition of Fgf7 or Fgf10 to the cultures induces formation of alveolus-like epithelial spheroids (‘alveolospheres’) comprised of intermingled AT1 and AT2 cells (see Brownfield et al. (*67*)) (Fig. 6, F and G). Inhibition of Notch signaling by addition of the γ-secretase inhibitor DAPT to the cultures increased the percentage of cells selected as AT2 cells from 58 ± 4% in controls to 87 ± 3% (n= 500 cells scored in 3 experiments). Similarly, blocking antibodies against Notch1 and Notch2 (*75*) increased AT2 cell selection to 72 ± 7%, and 88 ± 5%, respectively. These results support a role for Notch signaling mediated by Notch1 and Notch2 in alveolar epithelial patterning and lateral inhibition of AT2 fate (Fig. 6H). However, the culture experiments do not exclude the possibility that other (non-epithelial) sources could also contribute to alveolar epithelial patterning in vivo.

## Discussion

Our analysis of alveolar patterning and morphogenesis reveals a new mechanism by which alveoli form in the lung. In contrast to the classical (‘septation’) model in which the epithelium gives rise to large sacs that are later subdivided into alveoli by myofibroblasts, we show that alveoli form early and directly by budding and outgrowth of the epithelium from airway stalks between enwrapping smooth muscle cells (Fig. 7). These smooth muscle cells rearrange to become a ring of myofibroblasts at the alveolar entrance. This surprisingly simple and elegant way to form alveoli relies on precise spatial and temporal control of epithelial patterning and coordination with surrounding cells.

**Figure 7.**
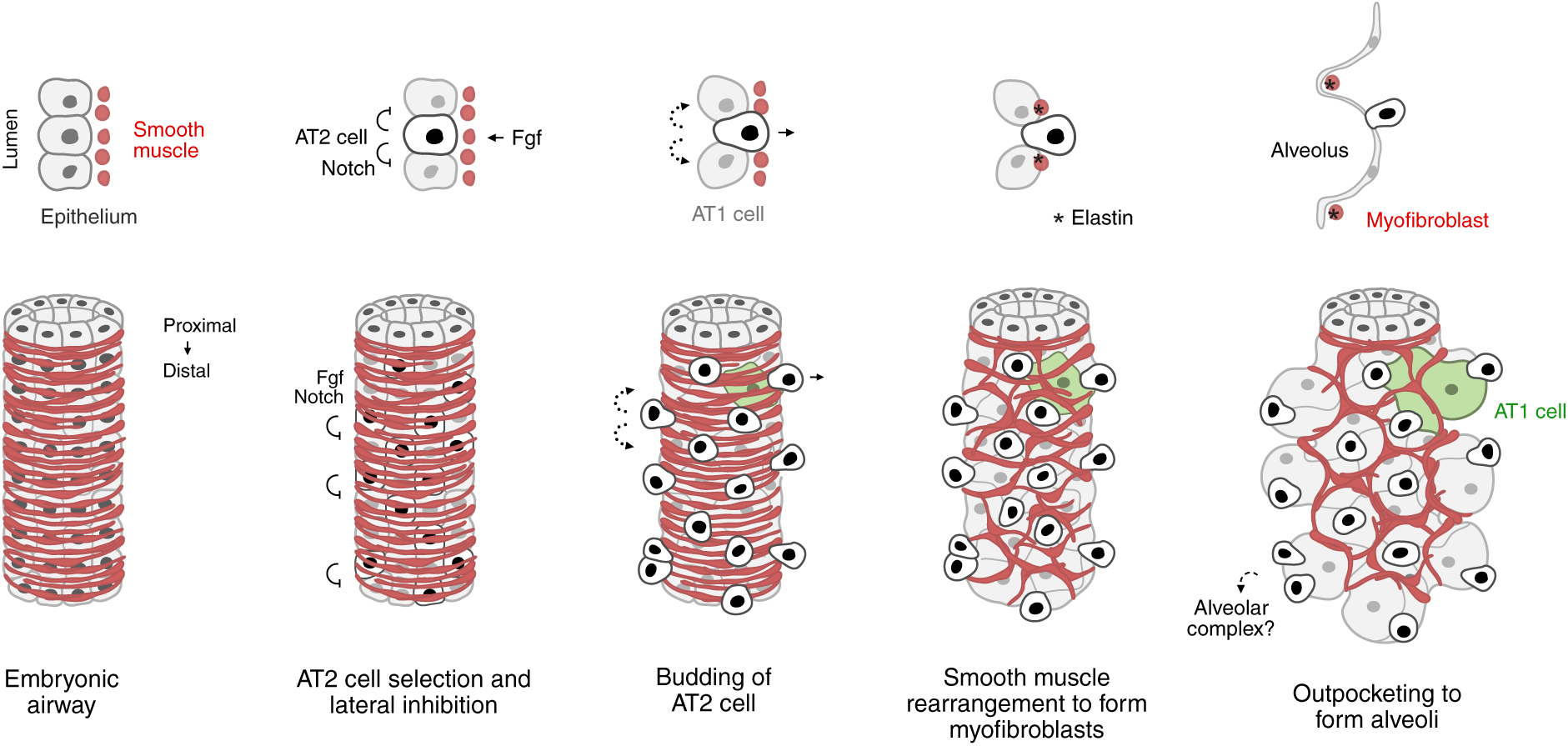
Model of alveolar patterning and formation by direct budding. Diagram of developing respiratory airway as side (top) and en face (bottom) views with epithelium (grey) and surrounding smooth muscle (red). Alveoli form on stalks by epithelial budding through the smooth muscle, led by single AT2 cells (black) that get selected by Fgf signaling and Notch-mediated lateral inhibition. Single AT2 cells are surrounded by 2-4 nascent AT1 cells (light grey) that are specified by another signal (dotted arrows), possibly a mechanical signal (*46*). As the nascent AT2 cell buds, smooth muscle cells rearrange around it, forming a ring of 3-5 myofibroblasts that deposit elastin (asterisk) and define the alveolar entrance. Groups of budding cells (2-3 cells, see Fig. 5B) may be sites of alveolar complex formation (dashed arrow). Outpocketing to form alveoli is accompanied by flattening and rounding of AT1 cells into a cup shape (one cell highlighted in green) and may be driven by mechanical forces around birth.

We propose that the initiating event in alveolar development is the selection of a single AT2 cell that first becomes apparent by Fgfr2 restriction in the embryonic lung (see also Brownfield et al., in review (*67*)) right after the boundary is set between conducting and respiratory airways (*57*). The nascent AT2 cell leads the alveolar outgrowth and is the source of one or more signals to surrounding cells: a lateral inhibition signal to neighboring epithelial cells mediated by Notch1 and Notch2 that prevents selection of more than one AT2 cell, and possibly a signal to nearby smooth muscle to rearrange to form entrance myofibroblasts and synthesize elastin. Our data support a mechanism of alveolar patterning analogous to Drosophila tracheal branching morphogenesis (*76*) where a single cell is selected by Fgf signaling to take the lead position in budding, and Notch signaling prevents selection of additional cells.

While Fgf and Notch signaling together control the selection and spacing of leading AT2 cells, and hence set the alveolar pattern, at least one other signal to the epithelium appears necessary to confer AT1 fate on surrounding cells and generate alveoli because expression of a constitutively-active Notch protein was insufficient to induce AT1 differentiation in cultured alveolar epithelial progenitors (not shown). A mechanical signal may play a role in AT1 specification and initial flattening, as suggested by a recent study (*46*), and pressure-induced forces may ensure further flattening and postnatal outgrowth of alveoli. Whatever the nature of this signal, it can be provided in cultures of purified alveolar epithelial cells induced with Fgf because such cultures readily form alveolospheres with intermingled AT1 and AT2 cells (Fig. 6F; see also Brownfield et al., in review (*67*)).

In our model of alveolar development, airway smooth muscle is the long sought source of entrance myofibroblasts. They synthesize elastin and are aligned with thick elastic fibers at the base of simple alveoli and alveolar complexes. Although the cells turn off smooth muscle actin in the postnatal (P12-15) lung, some and possibly all of them continue to be associated with alveolar openings – they may get anchored to the fibers or adhere to the basement membrane of AT1 or capillary cells through specialized junctional complexes (*34*). Genetic tools to specifically label and manipulate entrance myofibroblasts are needed to directly test the requirement of the cells in alveolus formation and to elucidate their role in the mature lung. Identification of the signals that recruit or activate them to make or repair fibers may suggest ways to prevent or reverse alveolar elastin destruction in age-associated and heritable forms of emphysema, which could lead to new strategies to replace damaged or missing alveolar entrance rings and inform treatments.

Our results also identified a second type of alveolar myofibroblast with a distinct anatomical location and developmental origin: ‘partitioning’ myofibroblasts that subdivide alveolar complexes. Interestingly, two molecular types of alveolar myofibroblasts were recently reported in a preprint investigating the diversity of myofibroblasts by scRNAseq at different stages of embryonic and postnatal lung development (*77*). One population expresses *Lgr6*, *Hhip* and *Cdh4* and persists post-alveologenesis, whereas the other expresses *Pdgfra* and undergoes developmental apoptosis. The *Lgr6*-expressing subset may correspond to entrance myofibroblasts that we show arise from the Lgr6 lineage, form the thick elastic fiber rings at the openings of simple and complex alveoli, and persist in the adult lung. The second population may be partitioning myofibroblasts that form thin fiber subdivisions of alveolar complexes.

The timing of alveolar complex formation and recruitment of partitioning myofibroblasts appears to coincide with the classical ‘alveolarization’ stage in the postnatal mouse lung (P4-P30) (*25, 27*). This suggests that these myofibroblasts and the associated thin fiber partitions form after the entrance rings of simple and complex alveoli and by a distinct mechanism, presumably part of septation. Although we limited our in depth analysis of alveolar development to stalks of early forming airways to avoid caveats associated with inflation of air-filled lungs and variability in late forming branches, it will be important to find ways around these biological and technical issues to probe at cellular resolution the origin and dynamics of partitioning myofibroblasts and the steps beyond entrance ring formation that expand and subdivide some alveoli into complexes (“septation”). Such approaches could also elucidate other steps in alveolar growth and maturation, including interlinking of alveoli through transseptal AT2 cells with multiple apical sides (Fig. S5C) and specialized circular junctions between AT1 cells from adjacent airways to form pores of Kohn (Fig. S5B).

Our model of alveolar development by direct budding is supported by the three-dimensional cellular structure of alveoli we elucidated by single-cell labeling. Remarkably, a simple alveolus in the mouse lung is composed of just 10-15 cells of 7 major cell types (excluding immune cells) including a single (‘lead’) AT2 cell, a single AT1 cell and 3-5 myofibroblasts at the entrance, which is 2-3 fold less than estimates from counting cells and alveoli on sections (*55*). Human alveoli are larger (200-300 µm mean diameter compared to 50 µm in mouse) and are thought to contain at least ten times more cells (*12, 16, 55, 78*). Single-cell labeling approaches should now be developed to elucidate the precise number and structure of cells in human alveoli and to determine if the mechanism of alveolar development we uncovered in mice, with selection of single lead AT2 cells as the key initiating and patterning event followed by budding and recruitment of entrance myofibroblasts from smooth muscle, is conserved between species. Based on our analysis, the timing of initiation of alveolar development in mouse and humans may be more similar than previously thought. And because humans also appear to have both simple alveoli and complexes (*2*), the two types of alveolar myofibroblasts, with their distinct origins, anatomy, and functions, may also be conserved. Our analysis of the structure and development of alveoli provides a basis for such evolutionary comparisons and for investigations of how the structure and cells of alveoli are altered with age and in human diseases that compromise gas exchange, including chronic lung diseases that arise in infants born too early with incompletely developed alveoli (*79*). By identifying the cells and molecules that control alveolar patterning and morphogenesis, our analysis suggests ways to stimulate failed or arrested alveolar development in premature lungs, and may inform strategies to rebuild alveoli in injured, aged or diseased lungs.

## Methods

### Mice

The following mouse strains were used: C57BL/6 (C57BL/6NCrl, Charles River Laboratories, strain 027) was the wild-type strain. *Hopx-CreER* (*Hopx^tm2.1(cre/ERT2)Joe^*/J) (*80*) (The Jackson Laboratory, strain 017606), *Foxa2-CreER* (*Foxa2^tm2.1(cre/Esr1*)Moon^*/J) (*81*) (The Jackson Laboratory, strain 008464), *Shh-GFP-Cre* (B6.Cg-*Shh^tm1(EGFP/cre)Cjt^*/J) (*82*) (The Jackson Laboratory, strain 005622), *VE-cadherin-CreER* (Tg(Cdh5-cre/ERT2)CIVE23Mlia) (*83*) (provided by M. L. Iruela-Arispe), *Apelin-CreER* (*Apln^tm1.1(cre/ERT2)Bzsh^*) (*84*) (provided by B. Zhou), *Aplnr-CreER* (*Tg(Aplnr-cre/ERT2)#Krh*) (*85*) (provided by K. Red-Horse), *Pdgfra-CreER* (B6N.Cg-Tg(Pdgfra-cre/ERT)467Dbe/J) (*86*) (The Jackson Laboratory, strain 018280), *Pdgfra-CreER* (B6.129S-*Pdgfra^tm1.1(cre/ERT2)Blh^*/J) (*87*) (The Jackson Laboratory, strain 032770), *Notch3-CreER* (*88*), *Gli1-CreER* (*Gli1^tm3(cre/ERT2)Alj^*/J) (*89*) (The Jackson Laboratory, strain 007913), *Lgr6-EGFP-IRES-CreERT2* (B6.129P2-*Lgr6^tm2.1(cre/ERT2)Cle^*/J) (*60*) (The Jackson Laboratory, strain 016934), *SMA-CreER* (Tg(Acta2-cre/ERT2)12Pcn) (*90*), and *SMMHC-CreER* (B6.FVB-Tg(Myh11-cre/ERT2)1Soff/J) (*64*) (The Jackson Laboratory, strain 019079) were used for expression of Cre recombinase. *Rosa26-mTmG* (B6.129(Cg)-*Gt(ROSA)26Sor^tm4(ACTB-tdTomato,- EGFP)Luo^*/J) (*91*) (The Jackson Laboratory, strain 007676), which expresses membrane targeted GFP after recombination, *Rosa26-tdTomato* (*Gt(ROSA)26Sor^tm14(CAG-tdTomato)Hze^*) (*61*) (The Jackson Laboratory, strain 007914), which expresses cytoplasmic tdTomato after recombination, *Rosa26-Rainbow* (Gt(ROSA)26Sor^tm1(CAG-EGFP,-mCerulean,-mOrange,-mCherry)Ilw^) (*65*) (provided by I. L. Weissman), which expresses cytoplasmic Cerulean, mOrange or mCherry after recombination, and *Rosa26-Confetti* (B6.129P2-*Gt(ROSA)26Sor^tm1(CAG-Brainbow2.1)Cle^*/J) (*92*) (The Jackson Laboratory, strain 017492), which expresses membrane targeted Cerulean CFP, nuclear GFP, cytoplasmic EYFP or cytoplasmic RFP after recombination, were used as Cre reporters. *CBF1-H2B-Venus* (Tg(Cp-HIST1H2BB/Venus)47Hadj/J) (*71*) (The Jackson Laboratory, strain 020942), which expresses nuclear Venus under the control of CBF1 binding sites, was used to visualize Notch activity in developing lungs. All experimental mice and embryos were heterozygous (or hemizygous) for indicated alleles. Only female mice and embryos were used for experiments with *Apelin-CreER*, as *Apelin* is X-linked (*84*). Noon of the day a vaginal plug was detected was considered as E0.5. The day a litter was born was considered as P0. For induction of Cre recombinase activity, tamoxifen (Sigma, Cat. No. T5648) was dissolved in corn oil and administered by intraperitoneal (i.p.) injection unless otherwise noted. Animal experiments were performed in accordance with approved Animal Care and Use Committee protocols.

### Immunostaining

Embryonic mouse lungs were dissected in PBS and fixed in 4% paraformaldehyde (PFA) for 1 hour (E16.5), 1.5 hours (E17.5) or 2 hours (E18.5) at 4°C. Fixed lungs were washed three times in PBS for 15 minutes at 4°C, then dehydrated through a PBS and methanol series into 100% methanol and stored at -20°C. For Sox2 antibody stains, embryonic lungs were fixed in 0.5% PFA for 3 hours at room temperature, followed by washes and dehydration into methanol. For Fgfr2 stains, lungs were incubated in Dent’s fixative (80% methanol, 20% dimethylsulfoxide (DMSO)) overnight at 4°C, washed twice in 100% methanol and stored as above.

Postnatal mouse lungs were perfused with PBS, inflated with 2% low-melting point agarose (Thermo Fisher, Cat. No. 16520100) dissolved in PBS and fixed by immersion in 4% PFA for 2 hours (P10) or 3-5 hours (adult) at 4°C (or in 0.5% PFA for 5 hours at room temperature for Sox2 antibody stains). Fixed tissue was washed three times in PBS for 30 minutes and dehydrated in methanol as above. Lungs were rehydrated on the day of staining through a methanol and PBS series into PBS. Sections (300-500 µm) were cut from left or right cranial lobes on a vibratome (Leica Biosystems). Rehydrated whole lobes or vibratome sections were blocked in 5% heat inactivated goat serum (or donkey serum for primaries raised in goat) in PBS with 0.5% Triton X-100 and 3% BSA for 2 hours at room temperature and incubated in primary antibodies diluted in block for 3 nights at 4°C.

Primary antibodies used, at indicated concentrations, were: rat anti-E-cadherin (1:500, Thermo Fisher, Cat. No. 13-1900, clone ECCD-2), goat anti-Sox2 (1:150, Santa Cruz, Cat. No. sc-17320), rabbit anti-Fgfr2 (1:150, Santa Cruz, Cat. No. sc-122, C-17), Armenian hamster anti-Muc1 (1:500, Thermo Fisher, Cat. No. HM1630P0, clone MH1), rabbit anti-SftpC (1:200, Millipore, Cat. No. AB3786), Syrian hamster anti-podoplanin (Pdpn; 1:50, DHSB, Cat. No. 8.1.1), rat anti-RAGE (1:500, R&D, Cat. No. MAB1179, clone 175410), Armenian hamster anti-Pecam1 (1:300, Bio-Rad, Cat. No. MCA1370Z, clone 2H8), rabbit anti-NG2 (1:200, Millipore, Cat. No. AB5320), rat anti-Pdgfrb (1:100, eBioscience, Cat. No. 14-1402), rabbit anti-elastin (1:500, kindly provided by Robert Mecham), goat anti-Itga8 (integrin α) (1:500, R&D, Cat. No. AF4076), goat anti-Pdgfra (1:200, R&D Systems, Cat. No. AF1062), rabbit anti-RFP (1:300, Rockland, Cat. No. 600-401-379), chicken anti-GFP (1:500, Abcam, Cat. No. ab13970), and mouse anti-SMA conjugated to FITC (1:200, Sigma, Cat. No. F3777, clone 1A4), Cy3 (1:200, Sigma, Cat. No. C6198, clone 1A4), or Cy5 (1:200). The SMA-Cy5 conjugate was prepared by mixing 100 µg of SMA antibody (Sigma, Cat. No. A2547, clone 1A4) with 4 µl of Cy5 NHS Ester (GE Healthcare; 1 mg dye dissolved in 10 µl of anhydrous DMSO) and incubation for 2 hours at room temperature in the dark. Unbound dye was removed using a gel filtration column (P-30, Bio-Rad) equilibrated in PBS with 0.09% sodium azide.

Following primary incubation, samples were washed in block six times for 1 hour and incubated in secondary antibodies conjugated to fluorescent dyes diluted 1:250 in block with added 4’,6-diamidino-2-phenylindole (DAPI; 1:500 dilution of 1 mg/ml stock, Thermo Fisher) for 2 nights at 4°C.

Secondary antibodies used, at indicated concentrations, were: goat anti-chicken Alexa Fluor 488 (Thermo Fisher, Cat. No. A-11039), goat anti-rabbit Alexa Fluor 568 (Thermo Fisher, Cat. No. A-11036), goat anti-rabbit Alexa Fluor 633 (Thermo Fisher, Cat. No. A-21070), goat anti-Armenian hamster Alexa Fluor 647 (Jackson Immunoresearch, Cat. No. 127-605-160), donkey anti-goat Alexa Fluor 488 (Thermo Fisher, Cat. No. A-11055), or donkey anti-rat Alexa Fluor 647 (Jackson Immunoresearch, Cat. No. 712-605-153).

Following secondary incubation, samples were washed in PBS with 0.1% Tween-20 six times for 1 hour, post-fixed in 4% PFA for 1 hour at 4°C, washed again three times for 15 minutes in PBS and dehydrated into methanol as above. Samples were cleared in benzyl alcohol:benzyl benzoate (BABB; 1:2) for confocal imaging.

For signal amplification of rat anti-GFP primaries, inflated and fixed whole lobes or vibratome sections were bleached with 5% hydrogen peroxide in methanol for 5 hours at room temperature, blocked as above and incubated in primary (rat anti-GFP, 1:200, ChromoTek, Cat. No. 3h9-100, clone 3H9) diluted in block as above. Samples were washed in block six times for 1 hour and incubated with horseradish peroxidase (HRP)-conjugated goat anti-rat IgG (Vector, Cat. No. PI-9401) diluted 1:150 in block for 2 nights at 4°C. Samples were washed again in PBS with 0.1% Tween-20 six times for 1 hour and incubated in fluorescein tyramide reagent (Perkin Elmer, Cat. No. SAT701001EA) diluted 1:70 in amplification diluent for 50 minutes at room temperature. Samples were washed in PBS four times for 15 minutes, post-fixed and cleared as above.

For visualizing elastic fibers, lungs were stained with fluorescent hydrazide (*50*). Whole embryonic or postnatal lobes, dissected and fixed as above in 4% PFA, were incubated with Alexa Fluor 350 hydrazide (Thermo Fisher, Cat. No. A-10439, 1:100 dilution of 2.5 mg/ml stock made up in PBS), Alexa Fluor 488 hydrazide (Cat. No. A-10436, 1:200 dilution of 0.5 mg/ml stock), Alexa Fluor 633 hydrazide (Cat. No. A-30634, 1:500 dilution of 0.5 mg/ml stock) or Alexa Fluor 647 hydrazide (Cat. No. A-20502, 1:200 dilution of 0.5 mg/ml stock) diluted in PBS with 0.1% Triton-X-100 for 1-3 nights at 4°C. Samples were washed in PBS with 0.1% Triton-X-100 four times for 30 minutes and cleared in BABB overnight or transferred to Cubic1 solution (*93*) right before imaging. Labeling of elastin by fluorescent hydrazide was verified by co-staining with rabbit anti-elastin antibodies (1:500, kindly provided by Robert Mecham) (see Fig. S2C). For co-stains with antibodies, fluorescent hydrazide was added during incubation with secondary antibodies.

### Single-molecule in situ hybridization (smFISH)

Postnatal lungs, inflated as described above, were fixed in 10% neutral buffered formalin (Fisher Scientific) for 24 hours at room temperature and transferred to 70% ethanol (made up in PBS) following 3 brief washes in PBS for embedding in paraffin. Sections were cut at 6 μm. smFISH was performed using a proprietary high-sensitivity RNA amplification and detection technology (RNAscope, Advanced Cell Diagnostics), according to the manufacturer’s instructions using the indicated proprietary probes, the RNAscope Multiplex Fluorescent Reagent Kit (v.2) and TSA Plus reagents (Perkin Elmer; 1:500 dilution for Cy3 and Cy5 dyes) or Opal dyes (Akoya Biosciences, 1:500 dilution for Opal 520). After smFISH, sections were incubated in DAPI (used at 2 μg/ml in PBS) for 5 min to counterstain nuclei and mounted in Prolong Gold antifade reagent (Invitrogen). Proprietary (Advanced Cell Diagnostics) probes used were: Mm-Fgf18 (495421), Mm-Eln (319361), Mm-Acta2-C2 (319531-C2), tdTomato (317041-C3).

### Mosaic labeling and quantification of alveolar cells

For labeling of individual AT1 cells, *Hopx-CreER* (*80*) or *Foxa2-CreER* (*81*) mice were crossed to *Rosa26-mTmG* (*91*) or *Rosa26-Confetti* (*92*) and dosed with 0.3-0.5 mg of tamoxifen 5 days prior to analysis at 2 months of age. Vibratome sections prepared from inflated and fixed lobes at 350 µm thickness were stained with chicken anti-GFP (to visualize membrane targeted GFP in *Rosa26-mTmG;* ChromoTek) or rabbit anti-RFP (to visualize cytoplasmic RFP in *Rosa26-Confetti;* Rockland) antibodies, Alexa Fluor 633 hydrazide and DAPI as described above, cleared and imaged in BABB. Confocal z-stacks were acquired using a 25x oil objective at an image resolution of 1,024 x 1,024 pixels and at a total thickness of 100-200 µm, which allowed visualization of entire alveoli since the average diameter of a mouse alveolus is 50-60 µm (*12*). Imaris Software (Bitplane) was used for 3D visualization of individual AT1 cells and their arrangement in alveoli and for surface reconstruction and quantification of the number of AT1 cells in an alveolus. The entrance to individual alveoli or complexes was identified as the ring formed by thick elastic fibers, visualized by hydrazide (see Fig. 1, A, C and E, Fig. S2-S4, Movies S1 and S2). Although the arrangement of individual AT1 cells was found to be variable, the surface covered by individual cells approximated the mean alveolar surface area (5,500 µm^2^) (*12*); hence the number of AT1 cells per alveolus is 1. Labeled AT1 cells in adult *Foxa2-CreER; Rosa26-Confetti* lungs were identified by immunostaining with rabbit anti-RFP (Rockland), Syrian hamster anti-Pdpn (DHSB) and rat anti-RAGE (R&D) antibodies (see Fig. S5A).

For mosaic labeling of capillary endothelial cells, *VE-cadherin-CreER* (*83*) (Fig. 1G and H) was bred to *Rosa26-Confetti* (*92*) and dosed with 1 mg of tamoxifen at 3 weeks. Lungs were analyzed at 2 months. The cell type was inferred from cell size and shape. For specific labeling of aerocytes and general capillary cells, *Apelin-CreER* (*84*) (Fig. S6A) or *Aplnr-CreER* (*85*) (Fig. S6B) mice were bred to *Rosa26-Confetti* and dosed with tamoxifen (0.5 mg for *Apelin-CreER*, 0.1 mg for *Aplnr-CreER*) at 2 months. Lungs were collected 5 days after tamoxifen injection. Vibratome sections (250 µm thickness) were stained with Alexa Fluor 633 hydrazide and endogenous RFP fluorescence was imaged in Cubic1-cleared sections, or stained with rabbit anti-RFP (Rockland) and Armenian hamster anti-Pecam1 (Bio-Rad) antibodies and fluorescent hydrazide as described above and cleared and imaged in BABB to visualize individual endothelial cells and their arrangement in the alveolar capillary network in Imaris. For quantification of the number of endothelial cells per alveolus, vibratome sections were stained with rabbit anti-Erg antibodies (1:100, Abcam, Cat. No. ab110639) to identify endothelial nuclei and fluorescent hydrazide to visualize elastic fibers. Endothelial nuclei and alveolar entrance rings were quantified in the Surpass view of Imaris.

For labeling of pericytes, *Pdgfra-CreER* (*86*); *Rosa26-mTmG* (Fig. 1I and Fig. S5E) or *Notch3-CreER* (*88*); *Rosa26-Confetti* lungs (Fig. S6C) were dosed with tamoxifen (2 mg for *Pdgfra-CreER*; 0.5 mg for *Notch3-CreER*) at 3 months. Lungs were collected at 5 days after tamoxifen injection and vibratome sections (350 µm) were stained with rat anti-GFP (to visualize membrane targeted GFP in *Rosa26-mTmG*; ChromoTek) or rabbit anti-RFP (to visualize cytoplasmic RFP in *Rosa26-Confetti*; Rockland) antibodies, rabbit anti-NG2 (Millipore) or rat anti-Pdgfrb (eBioscience) antibodies, and Alexa Fluor 633 hydrazide, cleared and imaged in BABB, and analyzed in Imaris. Pericytes were identified by NG2 or Pdgfrb co-staining. The transgenic *Pdgfra-CreER* strain labels multiple populations of lung mesenchyme including pericytes, as reported for the brain (*86*).

For labeling of alveolar fibroblasts, *Gli1-CreER* (*89*); *Rosa26-mTmG* (Fig. 1J and Fig. S5F) or *Pdgfra-CreER* (*87*)*; Rosa26-Confetti* lungs (Fig. S6D) were dosed with tamoxifen (4 mg for *Gli1-CreER*; 0.5 mg for *Pdgfra-CreER*) at 2 months. Lungs were collected at 5 days after tamoxifen injection and vibratome sections (350 µm thickness) were stained with rat anti-GFP and goat anti-Itga8 (Integrin α8) antibodies (R&D) to identify alveolar fibroblasts (*53, 62*) and with Alexa Fluor 633 hydrazide, cleared and imaged in BABB, and analyzed in Imaris.

For multicolor labeling of myofibroblasts, *SMMHC-CreER* (*64*) mice were bred to the multicolor Cre reporter *Rosa26-Rainbow* (*65*) and dosed with 0.1 mg tamoxifen at P5. Lungs were collected and stained with Alexa Fluor 647 hydrazide at P10, and endogenous Cerulean, mCherry and mOrange fluorescence was imaged in vibratome sections (250 µm thickness) cleared with ScaleA2 (incubation for 1-3 days at 4°C) (*94*). For quantification of myofibroblasts, vibratome sections from P10 *Lgr6-CreER* (*60*)*; Rosa26-tdTomato* lungs were stained with rabbit anti-RFP (Rockland) and mouse anti-SMA antibodies (Sigma) and fluorescent hydrazide, cleared and imaged in BABB, and analyzed in Imaris. Each intersection of neighboring alveoli was found to be occupied by a myofibroblast (identified by the lineage tag, expression of SMA and alignment with elastic fibers) and the cells form a ring around the alveolar entrance with no gaps. Each alveolus has 3-5 alveolar neighbors and is surrounded by 3-5 myofibroblasts.

For labeling of AT2 cells, *Shh-GFP-Cre* (*82*) mice were bred to *Rosa26-mTmG* (*91*) and analyzed at 2 months by immunostaining with chicken anti-GFP antibodies (Abcam) and Alexa Fluor 647 hydrazide, cleared and imaged in BABB. Clonal labeling was not necessary for AT2 cells because of their simple, well defined shape. For quantification of the number of AT2 cells per alveolus, vibratome sections (500 µm) from wild-type (C57BL/6) lungs were stained with rabbit anti-SftpC (Millipore) and Armenian hamster anti-Muc1 (Thermo Fisher) antibodies (to visualize AT2 cells), fluorescent hydrazide (to visualize elastic fibers and count alveolar units), and DAPI (to identify nuclei) as described above, and cleared and imaged in BABB. Confocal z-stacks taken were taken at an optical section thickness of 1 µm and a total thickness of 150 µm with a 25x oil objective, and the number of AT2 cells and gas exchange units (simple alveoli or individual units of alveolar complexes) were quantified in z-stacks (n=3 mice, one z-stack per mouse) in the Surpass view of Imaris. The number of AT2 cells (SftpC+ Muc1+ cells) was counted using the Imaris spot detection tool (stack 1: 338 AT2 cells, stack 2: 297 AT2 cells, stack 3: 370 AT2 cells) and the number of gas exchange units (the sum of the number of thick fiber rings at the entrance to simple alveoli and alveolar complexes and the number of thin fibers that subdivide alveolar complexes) was counted using the Imaris measurement point tool (stack 1: 300 gas exchange units, stack 2: 284 units, stack 3: 294 units). The average number of AT2 cells per gas exchange unit was calculated from these measurements (mean±s.d., 1.14±0.1 AT2 cells).

For quantification of the proportion of simple alveoli and alveolar complexes, 3 month-old C56BL/6 lungs (n=3) were stained with fluorescent hydrazide and z-stacks were taken at a thickness of 300 µm with a 25x oil objective. The elastic fiber network was visualized in three dimensions in the Surpass view of Imaris. The entrance to individual alveoli and complexes was identified as the ring formed by thick elastic fibers, visualized by hydrazide. Simple alveoli were scored as units that contain only thick fibers. Alveolar complexes were scored as units that also contain thin elastic fibers anchored on thick fiber rings. The number of thick fiber rings with and without thin fibers was quantified in z-stacks (n=3), corresponding to the number of alveolar complexes and simple alveoli, respectively.

For analysis of the structure of alveolar complexes, the number of alveolar epithelial cells was scored in adult *Foxa2-CreER; Rosa26-mTmG* lungs at 3 months stained with rabbit anti-SftpC (Millipore) and rat anti-RAGE (R&D) antibodies to visualize AT2 and AT1 cells (see Fig. S5A), respectively, fluorescent hydrazide to visualize elastic fibers and Dapi to identify nuclei. The number of AT1 and AT2 cells was scored in 2-unit (n=7) and 4-unit complexes (n=4) from 3D renderings of confocal z-stacks in the Surpass view of Imaris by counting alveolar epithelial nuclei.

### Mapping respiratory airways and alveolar development

The boundary between conducting and respiratory airways was mapped on the L.L1.A3 branch lineage (*56*). L.L1.A3 daughters are located at the flat edge of the left lobe (see Fig. S7A), which allowed imaging of whole lobes by confocal microscopy without sectioning. Conducting airways were defined as the generations of airways lined by a pseudostratified or columnar epithelium made up of Sox2-expressing airway epithelial cells. Respiratory airways were defined as the generations of airways lined by an epithelium made up of Sox2-negative alveolar epithelial cells (AT1, AT2 cells). The first-order respiratory airway (transitional bronchiole) was identified as a hybrid tube composed of a conducting and a respiratory part (*1*). In mice, the transition from conducting to respiratory airways is abrupt (*57*). The compartment boundary was identified by Sox2 immunostaining (*57*) as described above or by fluorescent streptavidin staining (*58*) to detect biotin expressed by airway epithelial cells: whole left lobes, dissected and fixed as described above in 4% PFA, were incubated in fluorescent streptavidin (Alexa Fluor 568 conjugate, Thermo Fisher, Cat. No. S-11226) diluted 1:200 in block (5% goat serum with 0.5% Triton X-100 and 3% BSA in PBS) for 3 nights at 4°C. Samples were washed in PBS with 0.1% Triton-X-100 five times for 1 hour and cleared and imaged in BABB. For co-stains with antibodies, fluorescent streptavidin was added during incubation with primary antibodies.

Alveolar development was mapped on respiratory airways of the first 2-3 generations (L.L1.A3 lineage), since branching in early generations is less variable and these airways form in the embryonic lung (*56*), which allows analysis of alveolar development without inflation. We analyzed stalks (rather than branch tips) to map the position of epithelial cells and smooth muscle cells relative to the axis of the airway tube.

### Lineage tracing of airway smooth muscle

*Lgr6-EGFP-IRES-CreERT2 (Lgr6-CreER)* (*60*) males were bred to *Rosa26-tdTomato* (*61*) females to generate heterozygous *Lgr6-CreER* males homozygous for *Rosa26-tdTomato*. The males were bred to CD1 wild-type females (Charles River Laboratories), since they were found to tolerate higher doses of tamoxifen than C57BL/6. Pregnant females from these crosses were dosed at E14.5 with saturating (4 mg) tamoxifen to label smooth muscle on embryonic airways with heritable tdTomato expression and the labeling was analyzed at E16.5, E17.5, P10, and P60 (Fig. S10E) by staining whole left lobes dissected and fixed as described above with rabbit anti-RFP (Rockland) and mouse anti-SMA (Sigma) antibodies, fluorescent streptavidin and hydrazide as described above, cleared in BABB and imaged on a Zeiss 880 confocal laser scanning microscope using a long working distance objective (LD Plan-Apochromat 20x/1.0 Corr M32 85mm).

### Clonal analysis of smooth muscle

*SMMHC-CreER* (*64*) mice were crossed to the multicolor Cre reporter *Rosa26-Rainbow* (*65*). Pregnant (E15) females received intraperitoneal injections of limiting doses (0.05 mg) of tamoxifen dissolved in corn oil to induce rare recombination events, which generated well-separated clones of cells that express one of three fluorescent proteins.

Postnatal (P10) lungs were perfused with PBS, inflated with 2% low-melting point agarose and fixed in 4% PFA for 2 hours at 4°C. Vibratome sections were prepared from left or right cranial lobes at 200-300 µm thickness. Sections were incubated with Alexa Fluor 350 hydrazide and SMA-Cy5 in PBS with 5% goat serum and 0.5% Triton X-100 for 2 nights at 4°C, washed in PBS with 0.1% Tween-20 five times for 15-30 minutes at 4°C and transferred to Cubic1 solution right before imaging. Endogenous Cerulean, mCherry and mOrange fluorescence was imaged on a Zeiss 880 confocal laser scanning microscope equipped with a 440 nm laser to excite Cerulean.

### Analysis of Notch activity in alveolar epithelial progenitors

Embryonic (E16.5-E18.5) *CBF1:H2B-Venus* (*71*) lungs were dissected and fixed in 4% PFA for 1-2 hours. Whole lobes were immunostained as above with chicken anti-GFP (Abcam), rabbit anti-Fgfr2 (Santa Cruz), rat anti-E-cadherin (Thermo Fisher) and Armenian hamster anti-Pecam1 (Bio-Rad) antibodies, cleared and imaged in BABB.

For Hes1 antibody stains, lungs were fixed in 0.5% PFA for 3 hours at room temperature, washed 3 times in PBS and incubated in 30% sucrose overnight at 4°C. Left lobes were embedded in optimal cutting temperature (OCT) compound and sectioned at 30 µm thickness. Cryosections were washed in PBS for 5 minutes and permeabilized in 0.3% Triton-X-100 for 10-15 minutes at room temperature. Sections were washed again in PBS for 5 minutes, blocked in 10% goat serum in PBS with 0.1% Tween-20 for 1-2 hours at room temperature, and incubated in primary antibodies diluted in block overnight at 4°C. Primary antibodies used, at indicated concentrations, were: rabbit anti-Hes1 (1:200, Cell Signaling, Cat. No. 11988), chicken anti-GFP (1:1000, Abcam, Cat. No. ab13970), Syrian hamster anti-podoplanin (Pdpn; 1:50, DHSB, Cat. No. 8.1.1), rat anti-RAGE (1:500, R&D Systems, Cat. No. MAB1179, clone 175410) and rat anti-E-cadherin (1:1000, Thermo Fisher, Cat. No. 13-1900). Sections were washed in PBS with 0.1% Tween-20 four times for 5 minutes at room temperature and incubated in secondary antibodies and DAPI (1:500 dilution of 1 mg/ml stock) diluted in block overnight at 4°C. Secondary antibodies used, at indicated concentrations, were: HRP-conjugated goat anti-rabbit IgG (1:150, Vector, Cat. No. PI-1000), goat anti-chicken Alexa Fluor 488 (1:500, Thermo Fisher, Cat. No. A-11039), goat anti-Syrian hamster Alexa Fluor 647 (1:500, Thermo Fisher, Cat. No. A-21451), donkey anti-rat Alexa Fluor 488 (1:500, Thermo Fisher, Cat. No. A-21208). Sections were washed again as above and incubated in Cy3 tyramide reagent (Perkin Elmer, Cat. No. SAT704A001EA) diluted 1:100 in amplification diluent for 10 minutes at room temperature. Sections were washed in PBS four times for 15 minutes, mounted in Prolong Gold antifade mountant (Thermo Fisher) and imaged on a Zeiss 780 confocal laser scanning microscope.

### Pharmacologic Notch inhibition in embryonic lungs

The γ-secretase inhibitor DBZ (Tocris, Cat. No. 4489) was delivered by daily intraperitoneal injection in timed-pregnant (E16-E18) C57BL/6 females at 30 µmol per kilogram body weight. A 100 mM DBZ stock was prepared in DMSO and diluted to a volume of 200 µl in corn oil. Controls were injected with an equivalent volume of DMSO diluted in corn oil. Lungs were dissected at E18.5 and fixed in 4% PFA for 2 hours at 4°C, immunostained for rabbit anti-SftpC (Millipore), Syrian hamster anti-Pdpn (DSHB) and rat anti-E-cadherin (Thermo Fisher) antibodies as described above, cleared and imaged in BABB on a Zeiss 780 confocal laser scanning microscope. 500 AT2 cells were scored for each condition (vehicle control, DBZ) in six biological replicates (n=6 mice per treatment group). For quantification of AT2 cell proliferation, lungs were immunostained with anti-SftpC and anti-Ki67-eFluor 570 antibodies (1:150, eBioscience, Cat. No. 41-5698-82) as above. AT2 cells (n=500) were scored for each condition (vehicle control, DBZ) in three biological replicates (n=3 mice per treatment group).

### Notch inhibition in cultured alveolar epithelial progenitors

Epithelial progenitors were isolated as previously described (*40*) from E16.5 C57BL/6 lungs pooled by litter and cultured on a layer of growth factor reduced Matrigel (BD) in serum-free media (DMEM/F12 supplemented with L-glutamine and penicillin-streptomycin) with added FGF ligands (Fgf7; 4 µM, R&D Systems, Cat. No. 5028-KG-025), the γ-secretase inhibitor DAPT (10 µM; Tocris, Cat. No. 2634), or Notch1 or Notch 2 antibody antagonists (*75*) (kindly provided by Chris Siebel, Genentech Inc.; 20 µg of antibody per ml of culture). Cultures were fixed after 4 days, immunostained for rabbit anti-SftpC (Millipore) and rat anti-RAGE antibodies (R&D) as described above to identify alveolar epithelial progenitors (SftpC+ RAGE+), AT1 cells (SftpC- RAGE+) and AT2 cells (SftpC+ RAGE-), and mounted in Vectashield media with DAPI (Vector) for imaging on a Zeiss 780 confocal laser scanning microscope. 500 cells were scored for each condition (Fgf7 only control, Fgf7 with DAPT, Fgf7 with anti-Notch1 antibody, Fgf7 with anti-Notch2 antibody) in three biological replicates (independent cultures established from 3 litters).

### Confocal imaging

Images were acquired using inverted Zeiss 780 or upright Zeiss 880 confocal laser scanning microscopes. Zen Imaging Software (Zeiss) and Adobe Photoshop were used to adjust image levels and pseudocolor the images. Volocity Software (Perkin Elmer) was used to generate maximum intensity projections from z-stacks. The fine filter was applied to images of Sox2 antibody stained lungs. Imaris Software (Bitplane) was used for 3D visualization.

### Analysis of single-cell RNA-sequencing (scRNAseq) data

Processed scRNAseq MARS-Seq data for developing (E16.5, E18.5) mouse lung by Cohen et al. (*74*) were obtained from GEO (accession number GSE119228) as gene count tables with de-multiplexed and aligned reads and expression profiles of cells were clustered using the R software package Seurat (version 2.3) (*95*). In brief, cells with fewer than 500 unique molecular identifiers were excluded and data were log-transformed: ln(unique molecular identifiers per ten thousand+1) [ln(UP10K+1)]. Highly variable genes were selected using the ‘FindVariableGenes’ function (dispersion (mean/variance) z-score > 0.5) for linear dimensionality reduction using principal component analysis. The number of principal components was selected by inspection of the plot of variance explained. Cells were clustered by constructing a shared nearest neighbor graph and clusters were visualized by t-distributed stochastic neighbor embedding (tSNE). Lung epithelial, endothelial, and stromal cells were identified by *Cdh1*, *Cdh5* or *Col1a1* expression, respectively. AT2 cells were identified by *Etv5*, *Sox9*, *Fgfr2*, *Lamp3* and *Sftpa1* expression. AT1 cells were identified by *Ager*, *Hopx*, *Pdpn* and *Aqp5* expression. Alveolar epithelial progenitor cells were identified by co-expression of AT2 (*Etv5*, *Sox9*, *Fgfr2*, *Lamp3*, *Sftpa1*) and AT1 markers (*Ager*, *Hopx*, *Pdpn*) (*40, 47*). Heatmaps were generated with the ‘heatmap.3’ function. Analysis of scRNAseq data for E18.5 mouse lung cells (Treutlein et al.) was conducted as previously described (*47*). Processed scRNAseq Smart-Seq2 data for adult mouse lung (Tabula Muris Senis) were obtained from Synapse (Synapse ID: syn21041850) (*62, 63*) as a Seurat object with annotated cell types. Dot plots were generated using the ‘DotPlot’ function in Seurat.

### Statistics and reproducibility

Data analysis and statistical tests were performed using R software (version 3.5.1). Data are represented as mean ± standard deviation (s.d.). For comparison of two groups, a two-sided Wilcoxon rank sum test was conducted at 5% significance level. Replicate experiments are biological replicates with different animals. For all graphs, the number of biologically independent samples is reported in the Figure Legends. Sample size calculations were not performed. Mice of the appropriate age were allocated into experimental groups (control vs. treatment) at random. The investigators were not blinded to sample allocation.

## Acknowledgements

We thank C. Siebel and R. Mecham for reagents; J. Campbell for help with culture experiments; Maya Kumar for advice and discussion; M. Petersen for help with illustrations; the Department of Comparative Medicine Animal Histology Services for technical assistance; M. L. Iruela-Arispe, K. Red-Horse, I. L. Weissman and B. Zhou for sharing mouse lines; and members of the Krasnow lab for comments on the manuscript. This work was supported by the Vera Moulton Wall Center for Pulmonary Vascular Disease at Stanford and grants from the Austrian Science Fund (J-3373) and the American Heart Association (16POST27250261) to A.G. K.J.T. was supported by a Paul and Mildred Berg Stanford Graduate Fellowship. M.A.K. is an investigator of the Howard Hughes Medical Institute.

## Author Contributions

A.G. and M.A.K. conceived, designed, and analyzed experiments and wrote the manuscript. A.G. performed the experiments. A.G. and K.R.St.J. performed lineage tracing. D.G.B. performed culture experiments. A.G. and D.G.B. analyzed scRNAseq data. A.G., K.J.T., and R.J.M performed mosaic labeling. R.J.M. provided guidance on the project. All authors reviewed the manuscript.

## List of Supplementary Materials

Figures S1-S17

Table S1. Cre and CreER drivers used for clonal labeling and lineage tracing.

Table S2. Airway clones generated by tamoxifen induction of *SMMHC-CreER; Rosa26-Rainbow*.

Movie S1. Three-dimensional architecture of a simple alveolus.

Movie S2. Three-dimensional architecture of an alveolar complex.

## Supplementary Figure Legends

**Figure S1.**
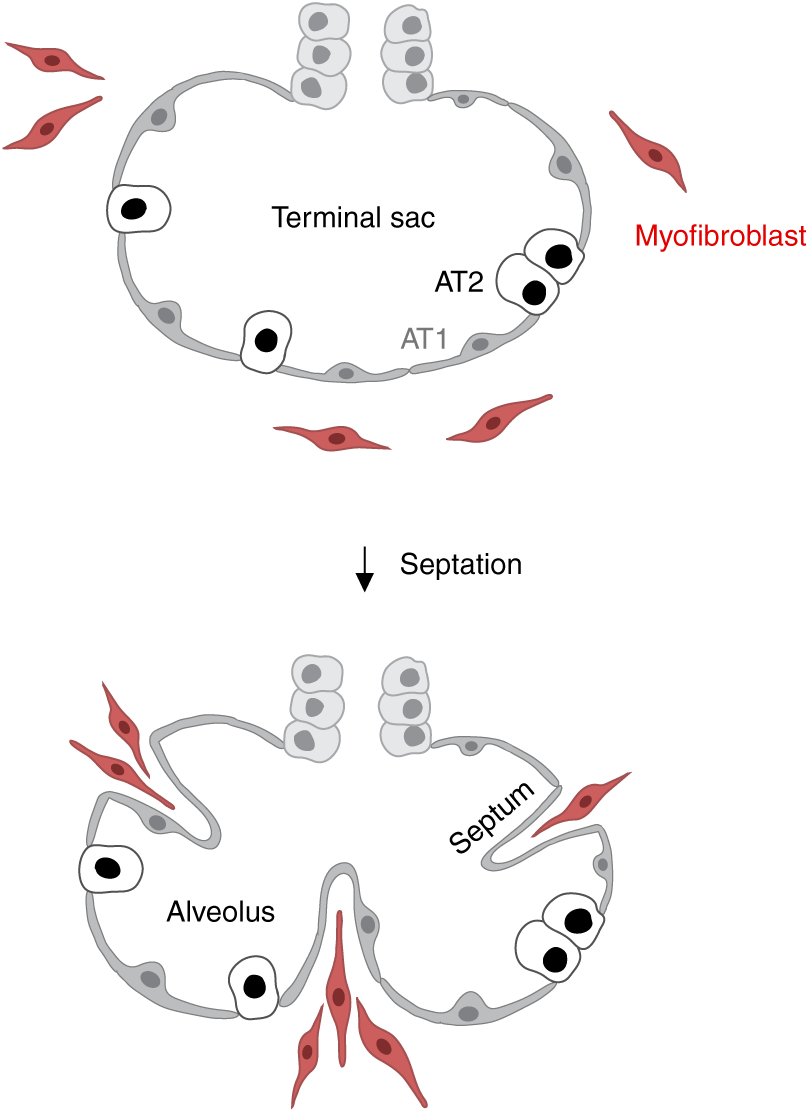
Textbook model of alveolar development by septation. Following branching morphogenesis (‘pseudoglandular’ stage, mouse: E9.5-E16.5, human: 5-17 weeks), the epithelium in distal branch tips differentiates to form squamous AT1 cells (grey) and cuboidal AT2 cells (black) (‘canalicular’ stage, mouse: E16.5-E17.5, human: 17-24 weeks) and the airspaces widen to form thin-walled saccules (‘saccular’ stage, mouse: E17.5-P4, human: 24 weeks to birth). The terminal sacs (top panel) are subdivided in the postnatal mouse lung into alveoli (bottom panel) by ingrowth of ridges or crests (‘secondary septa’) (‘alveolarization’ stage, mouse: P4-P30 (*25, 27*); human: 29-33 weeks to several years (*30, 31, 97*)). Septation is thought to be driven by contraction or inward migration of myofibroblasts (red) (*25-29, 44, 96*).

**Figure S2.**
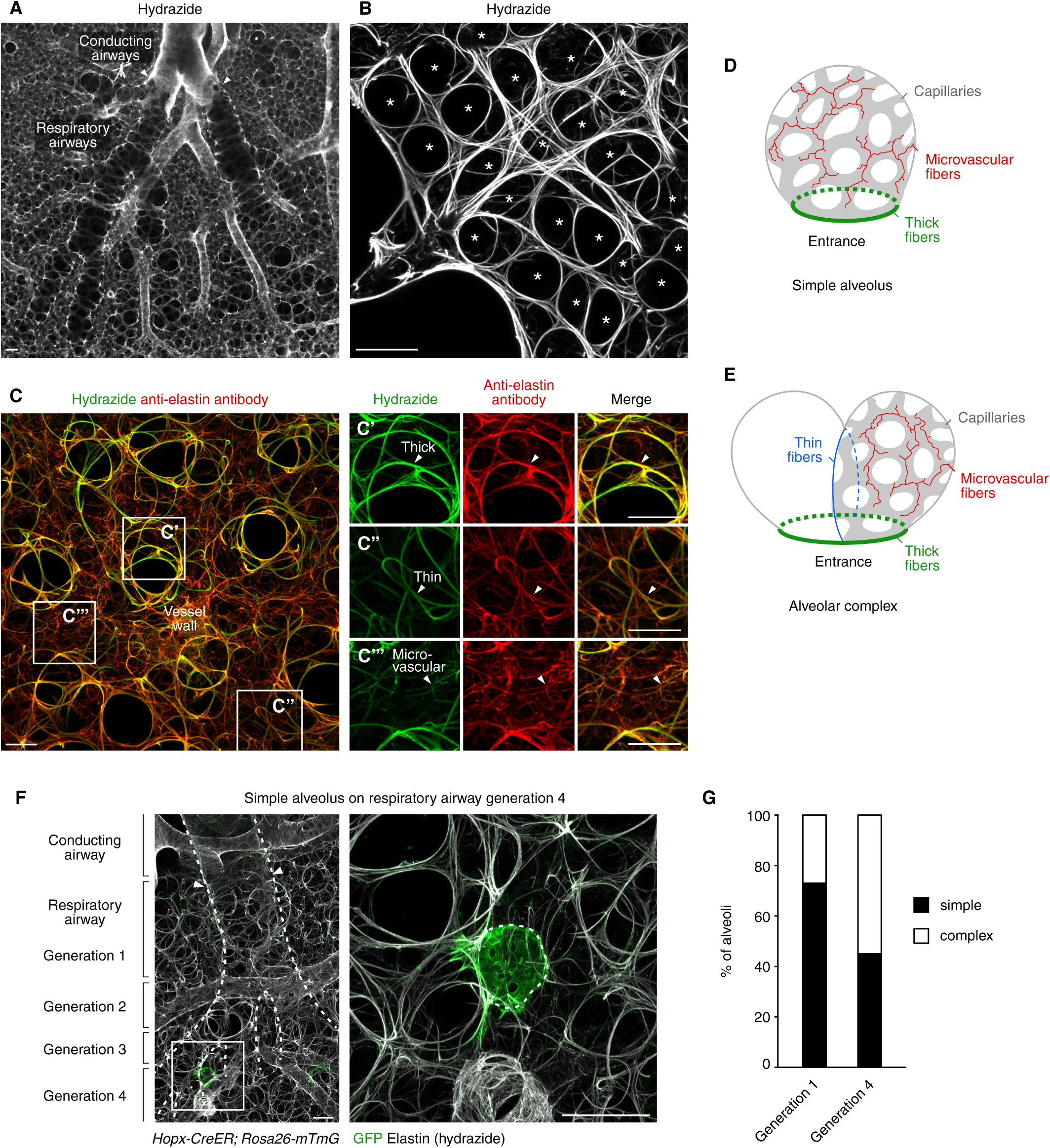
Structure and types of alveolar elastic fibers. (**A** and **B**) Confocal slice (**A**) and projection (**B**) showing conducting and respiratory airways (**A**) and alveolar entrance rings (**B**) in a 2 month-old wild-type lung stained with fluorescent hydrazide to visualize elastic fibers (white). Arrowheads, boundary between conducting and respiratory airways (note the abrupt transition in mice) (*57*). Scale bars, 50 µm. (**C**) Confocal projection with close-ups of boxed regions (**C’** to **C’’’**) showing three types of alveolar elastic fibers (thick, thin and microvascular fibers; arrowheads in **C’** to **C’’’**) in a 2 month-old lung stained with fluorescent hydrazide (green) and anti-elastin antibodies (red). Note correspondence between hydrazide and elastin antibody stains. Scale bars, 25 µm. (**D** and **E**) Diagrams showing the arrangement of elastic fibers in simple alveoli (**D**) and alveolar complexes (**E**). Thick fibers (green circles) form a ring at the alveolar entrance, microvascular fibers (red) are interwoven with capillaries (grey), and thin fibers (blue) subdivide a complex into individual units (the complex depicted in **E** is composed of 2 units). (**F**) Simple alveolus on late generation respiratory airway. Respiratory airway generations (left panel, dashed lines) are numbered (arrowheads, generation 1). Inset (right) shows close-up of boxed region highlighting a single, GFP-labeled AT1 cell (green) in this *Hopx-CreER; Rosa26-mTmG* lung dosed with limiting tamoxifen (0.3 mg) 5 days prior to analysis at 2 months of age and co-stained with GFP antibodies to visualize labeled AT1 cells and fluorescent hydrazide to label elastic fibers (white). (**G**) Quantification of the proportion (in % of alveoli) of simple alveoli and alveolar complexes in adult mouse lung (n=3 C57BL/6 lungs at 3 months of age stained with hydrazide; see Methods).

**Figures S3.**
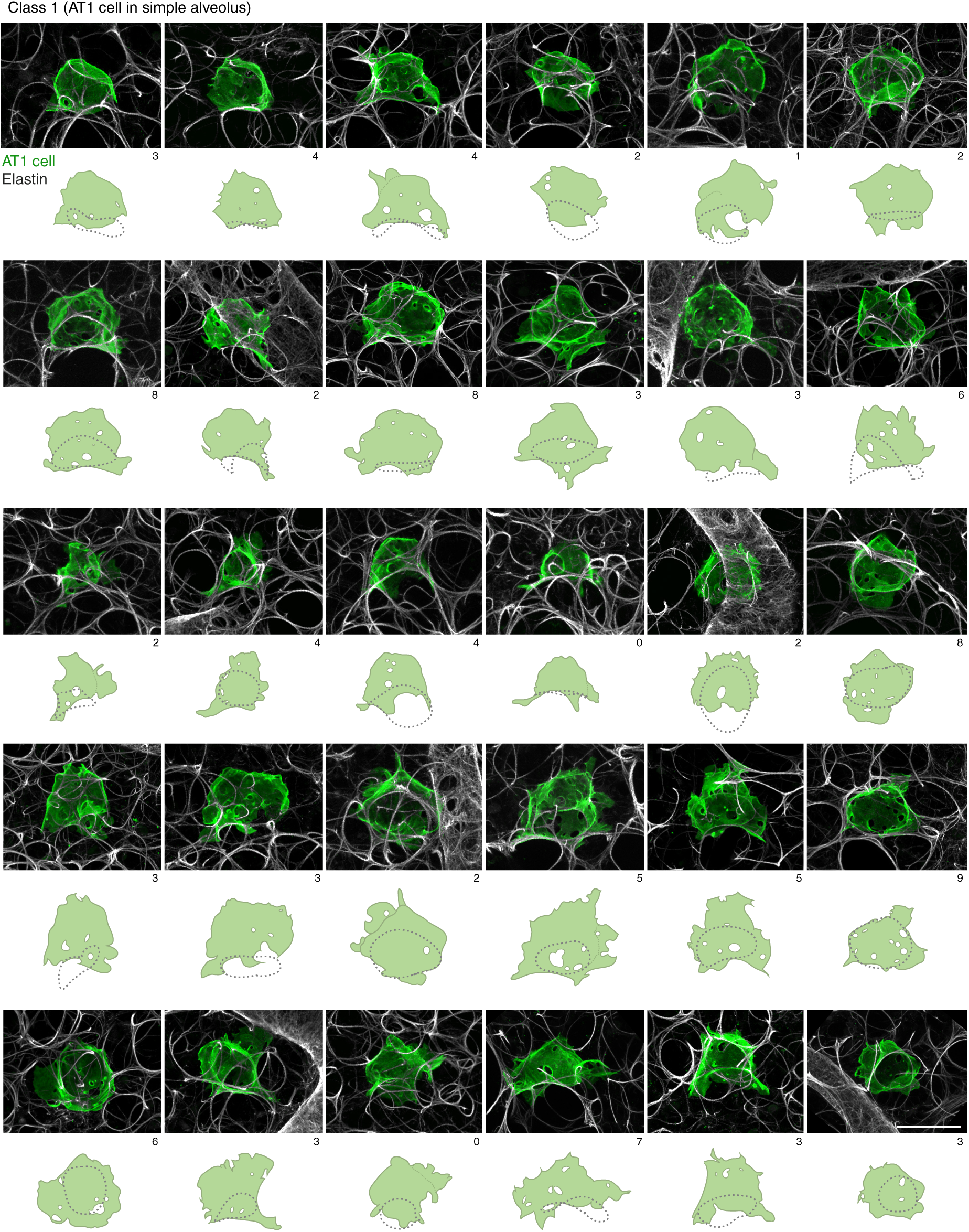
Morphology and arrangement of AT1 cells in simple alveoli. Confocal projections (top) and diagrams (bottom) showing the arrangement of single AT1 cells (green) in simple alveoli (class 1). Note pores of Kohn (membrane-lined ‘pores’ (channels) through AT1 cells (*52*) that connect the lumens of adjacent alveoli, white ovals in diagrams; the number of pores (0-9) per cell is indicated below projection images) and thick elastic fibers forming a ring at the alveolar entrance (dotted circles in diagrams). *Hopx-CreER* or *Foxa2-CreER; Rosa26-mTmG* lungs were dosed with tamoxifen (0.3 mg) 5 days prior to analysis and stained at 2 months of age with GFP antibodies and fluorescent hydrazide (elastic fibers, white). Scale bars, 50 µm.

**Figure S4.**
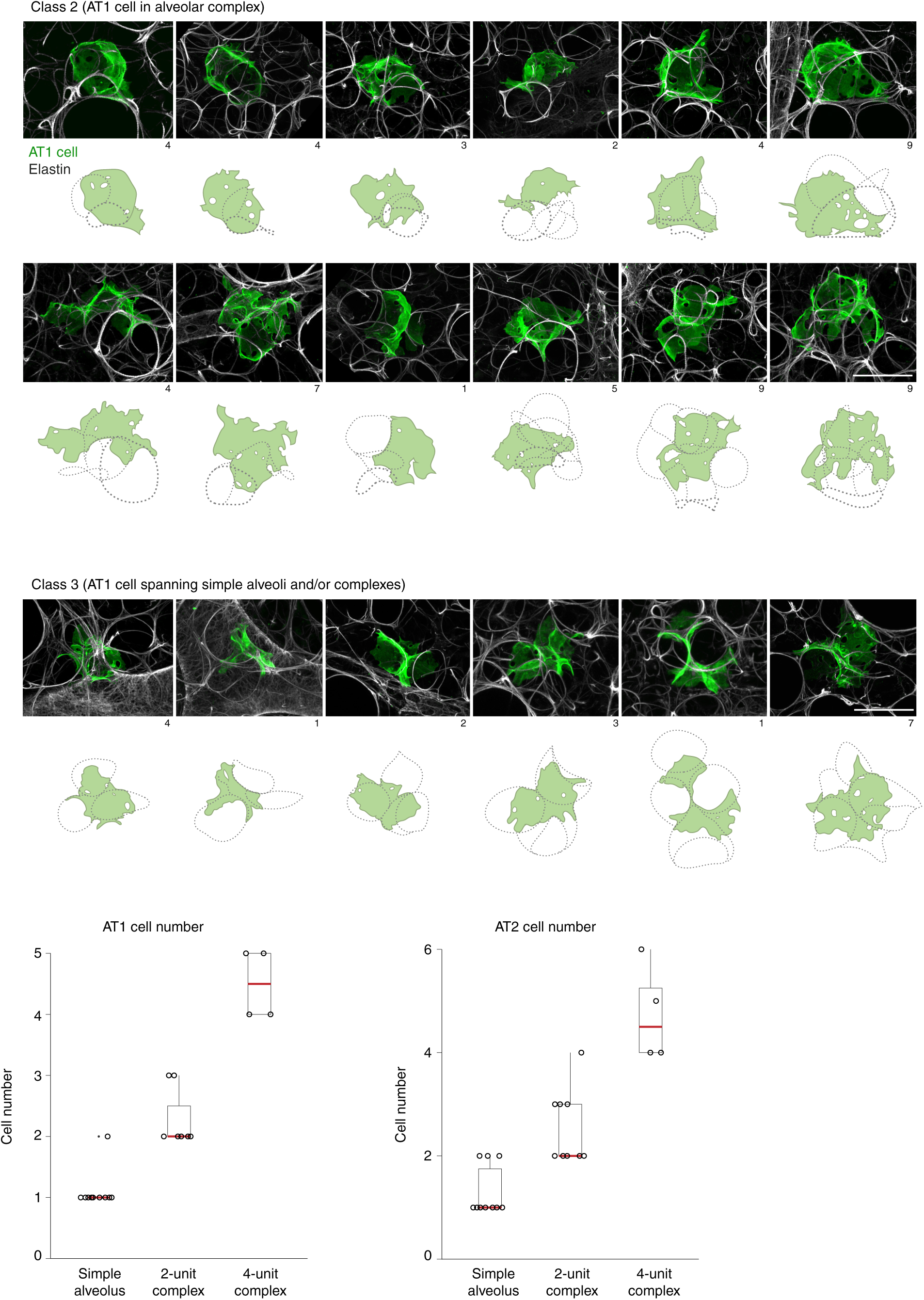
Morphology and arrangement of complex AT1 cells. Confocal projections (top) and diagrams (bottom) as in Figure S3 showing the arrangement of single AT1 cells (green) as part of alveolar complexes (class 2). Note pores of Kohn (white ovals in diagrams; the number of pores is indicated below projection images), thick elastic fibers forming a ring at the entrance of the alveolar complex (thick dotted circles) and thin fibers that subdivide alveolar complexes (thin dotted lines originating and terminating on thick dotted circle). Some AT1 cells could not be assigned to class 1 or class 2, since they were positioned between alveoli, and they may be part of simple alveoli and/or alveolar complexes (class 3). Scale bars, 50 µm. Bottom graphs show quantification of the number of AT1 (left) and AT2 cells (right) in simple alveoli and alveolar complexes. Data are shown as boxplots with individual data points plotted. The number of cells was scored from 3D renderings of confocal z-stacks as described in the Methods. AT1 cells scored in n=10 simple alveoli and n=11 alveolar complexes; AT2 cells scored in n=10 simple alveoli and n=13 alveolar complexes in adult mouse lung (n=3 lungs at 3 months of age stained with hydrazide; see Methods). Red bar, median value.

**Figure S5.**
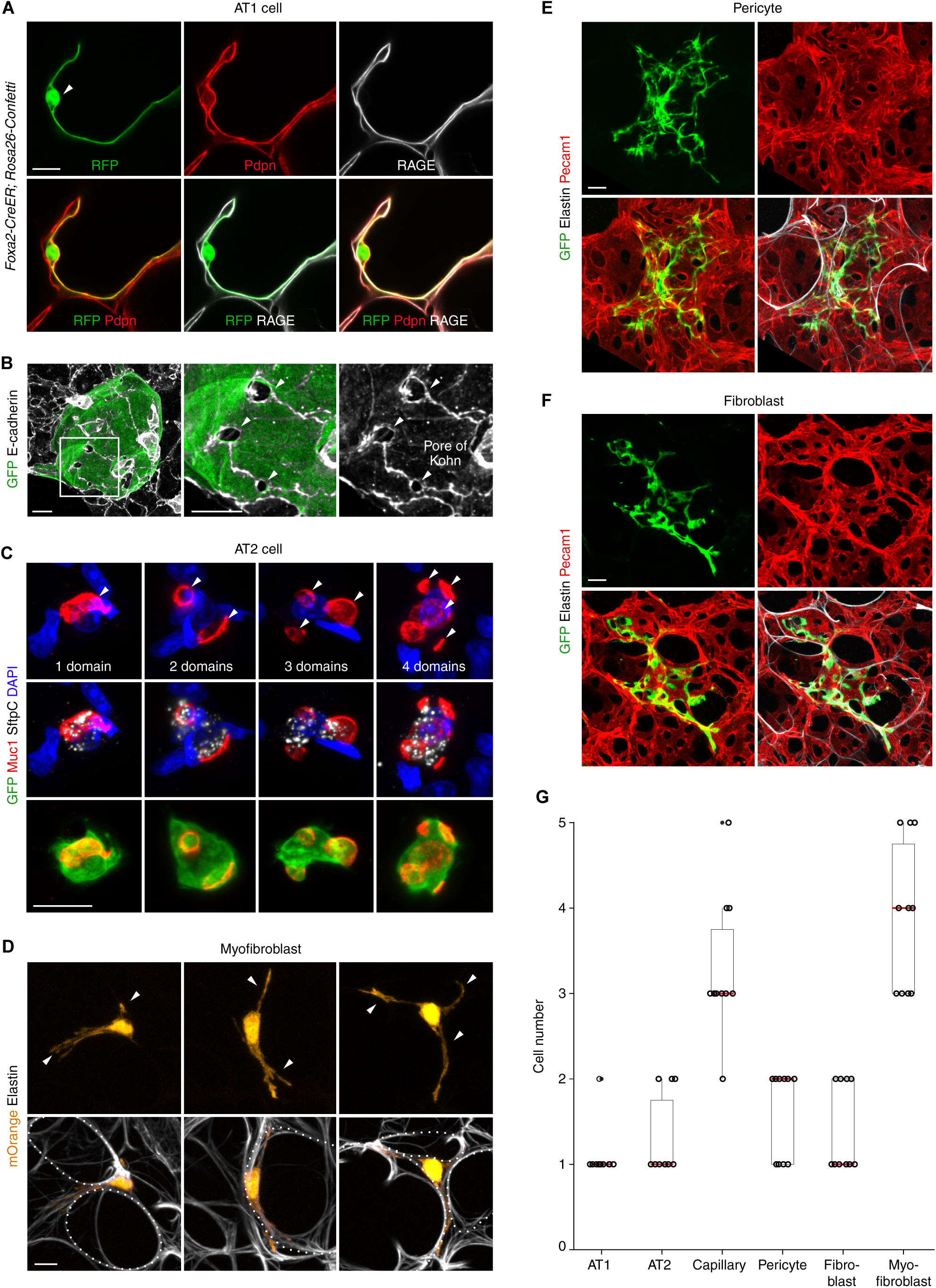
Morphologies and features of alveolar cells. (**A**) Confocal slice of a single AT1 cell (green) in a *Foxa2-CreER; Rosa26-Confetti* lung dosed with 0.5 mg tamoxifen and immunostained at 2 months for RFP (pseudocolored in green), the apical AT1 marker Pdpn (red) and the basal AT1 marker RAGE (white). Adult AT1 cells co-express Pdpn and RAGE and have an expansive, squamous shape. Arrowhead, AT1 cell nucleus. Scale bar, 10 µm. (**B**) Confocal projection of a single AT1 cell (green) with close-up of boxed region showing pores of Kohn (arrowheads) in a *Hopx-CreER; Rosa26-mTmG lung* dosed with 0.3 mg tamoxifen and stained at 2 months for GFP (green) and E-cadherin (white). Pores are surrounded by circular junctions formed by AT1 cells from adjacent airways. Individual pores are also connected by junctions (presumably autocellular junctions formed by a single AT1 cell). Scale bars, 10 µm. (**C**) Confocal projections of single AT2 cells (green) with 1-4 apical domains (red, arrowheads) in a *Foxa2-CreER; Rosa26-mTmG* lung dosed with 0.3 mg tamoxifen and stained for GFP (green), Muc1 (apical domains, red), SftpC (surfactant, white) and DAPI (nuclei, blue) at 2 months of age. Scale bar, 10 µm. (**D**) Confocal projections of single myofibroblasts (orange) that are aligned with thick elastic fibers (white) in a postnatal day 10 *SMMHC-CreER; Rosa26-Rainbow* lung dosed with 0.1 mg tamoxifen at P5 and stained with fluorescent hydrazide (elastin, white). Myofibroblasts have a simple spindle shape with 2 or 3 cell processes (arrowheads). Dotted lines, entrance to simple alveoli or complexes. Scale bar, 10 µm. (**E**) Confocal projection of pericyte (see also Fig. 1I) in a *Pdgfra-CreER; Rosa26-mTmG* lung dosed with 2 mg tamoxifen at 3 months and immunostained for GFP (green), Pecam1 (red) and elastin (visualized by hydrazide, white). Scale bar, 10 µm. (**F**) Confocal projection of alveolar fibroblast (see also Fig. 1J) in a *Gli1-CreER; Rosa26-mTmG* lung dosed with 4 mg tamoxifen at 2 months and immunostained for GFP (green), Pecam1 (red) and elastin (visualized by hydrazide, white). Scale bar, 10 µm. (**G**) Quantification of the number of cells in a simple alveolus. Data are shown as boxplots with individual data points plotted; n=10 cells of each type scored in adult mouse lungs (n=3 lungs at 3 months of age) with mosaic labeling and immunostaining against common antigens as described in the Methods. Red bar, median value.

**Figure S6.**
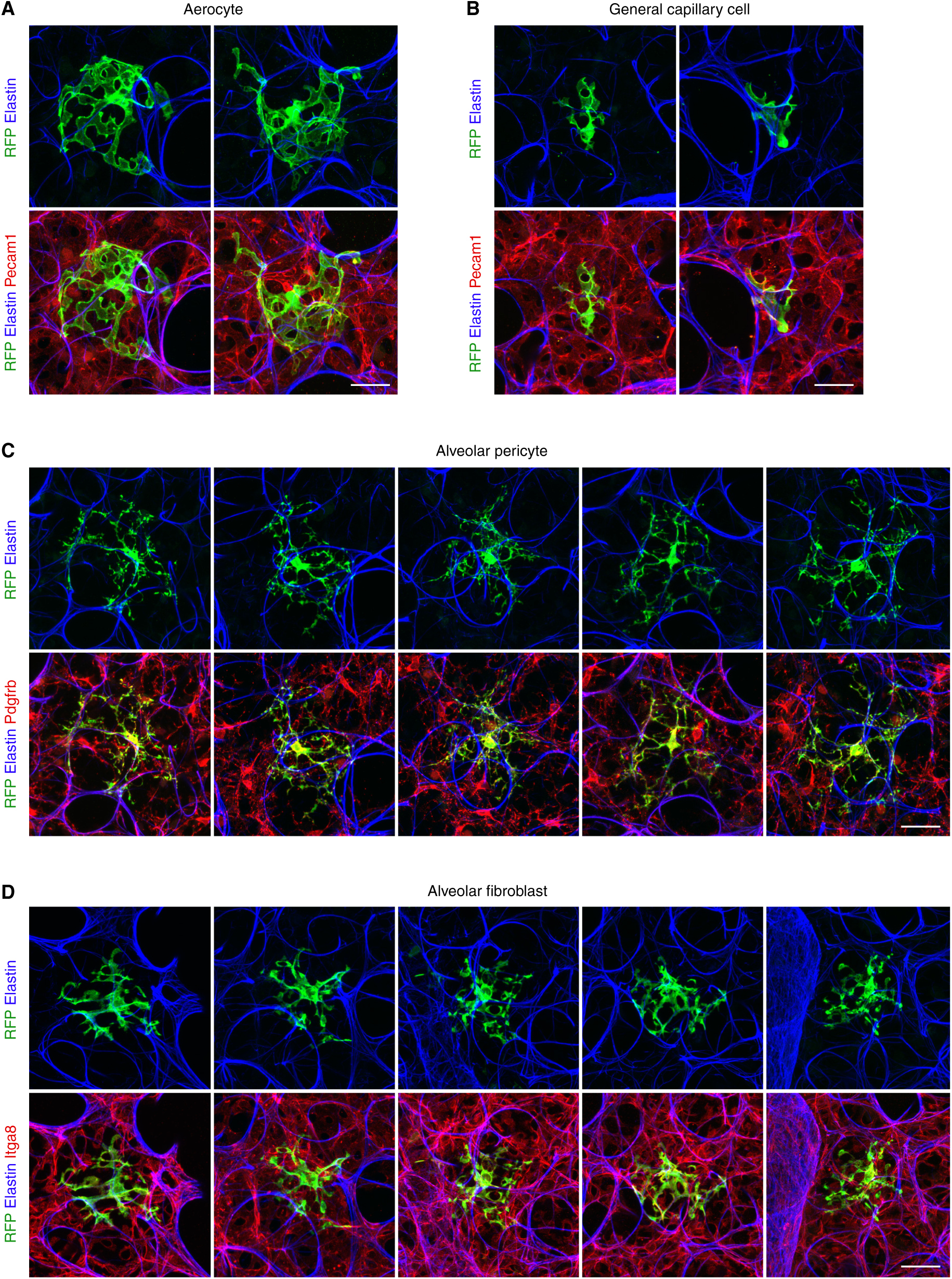
Single cell labeling of alveolar cells. (**A**, **B**) Confocal projections of single aerocytes in *Apelin-CreER; Rosa26-Confetti* lungs (**A**) and single general capillary cells in *Aplnr-CreER; Rosa26-Confetti* lungs (**B**) dosed with tamoxifen (**A**, 0.5 mg; **B**, 0.1 mg) at 2 months and immunostained for RFP (pseudocolored in green), Pecam1 (red) and elastin (visualized by hydrazide, blue) (see also Gillich et al. (*51*)). Scale bar, 10 µm. (**C**) Confocal projections of single pericytes in *Notch3-CreER; Rosa26-Confetti* lungs dosed with 0.5 mg tamoxifen at 3 months and immunostained for RFP (pseudocolored in green), Pdgfrb (red) and elastin (visualized by hydrazide, blue). Scale bar, 10 µm. (**D**) Confocal projections of single alveolar fibroblasts in *Pdgfra-CreER; Rosa26-Confetti* lungs dosed with 0.5 mg tamoxifen at 2 months and immunostained for RFP (pseudocolored in green), Integrin α8 (Itga8, red) and elastin (visualized by hydrazide, blue). Scale bar, 10 µm.

**Figure S7.**
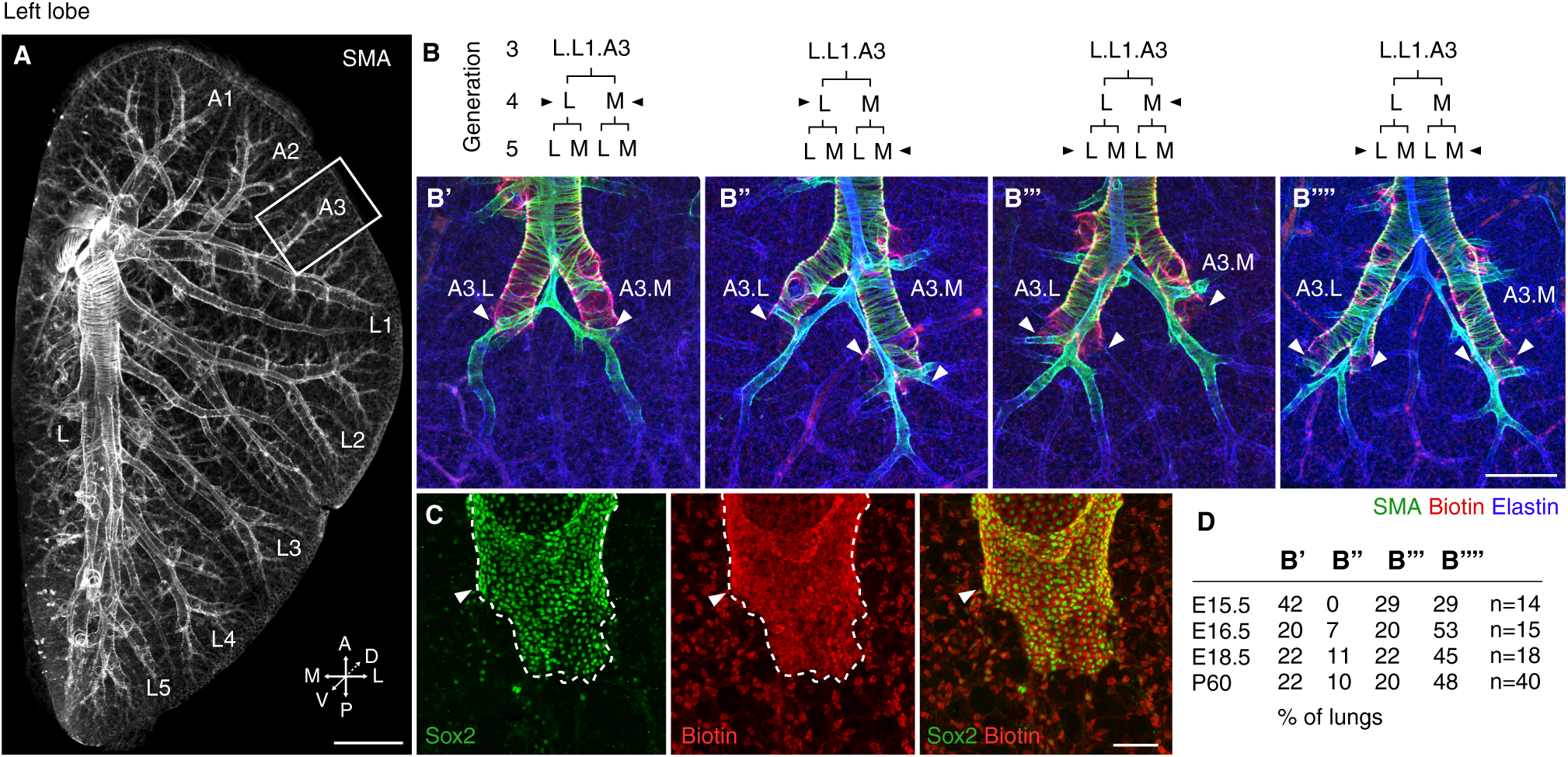
Mapping the boundary between conducting and respiratory airways. (**A**) Left lobe of a wild-type (C57BL/6NCrl) lung immunostained for SMA (smooth muscle, white) with boxed region showing L.L1.A3 branch and its daughters (see also lineage diagram in Fig. 2A). M, medial, L, lateral, A, anterior, P, posterior, V, ventral, D, dorsal. Scale bar, 500 µm. (**B**) Lineage diagrams (top) and confocal projections (bottom) showing the position of the boundary between conducting and respiratory airways (arrowheads) on L.L1.A3 daughter branches in wild-type lungs stained at 2 months of age with SMA antibodies (smooth muscle, green), fluorescent streptavidin to detect biotin expressed by airway epithelial cells (red) and fluorescent hydrazide (elastin, blue). Panels show four examples of boundary patterns observed along L.L1.A3: symmetric at generation 4 (**B’**), asymmetric at generation 4 on the lateral aspect of L.L1.A3 and at generation 5 on the medial side (**B’’**), asymmetric at generation 5 on the lateral side and generation 4 on the medial side (**B’’’**), and symmetric at generation 5 (**B’’’’**). See quantification in panel **D**. Scale bar, 500 µm. (**C**) Confocal projections showing that the compartment boundary (arrowhead in green panel) visualized by Sox2 expression (green) corresponds to the biotin expression domain (arrowhead in red panel) detected by fluorescent streptavidin (red) in a 2 month old wild-type lung stained with anti-Sox2 antibodies and Alexa Fluor 568 streptavidin. Scale bar, 50 µm. (**D**) Table showing the percentage of wild-type (C57BL/6NCrl) lungs at the indicated embryonic and adult ages with the boundary patterns shown in **B’** to **B’’’’**. Note that the percentage of lungs with pattern B’ (symmetrical at generation 4) is similar at E16.5, E18.5, and P60 (20-22%), but higher in E15.5 (42%), indicating that the junction is dynamic until it is permanently established at around E16-E16.5, consistent with previous studies (*57*).

**Figure S8.**
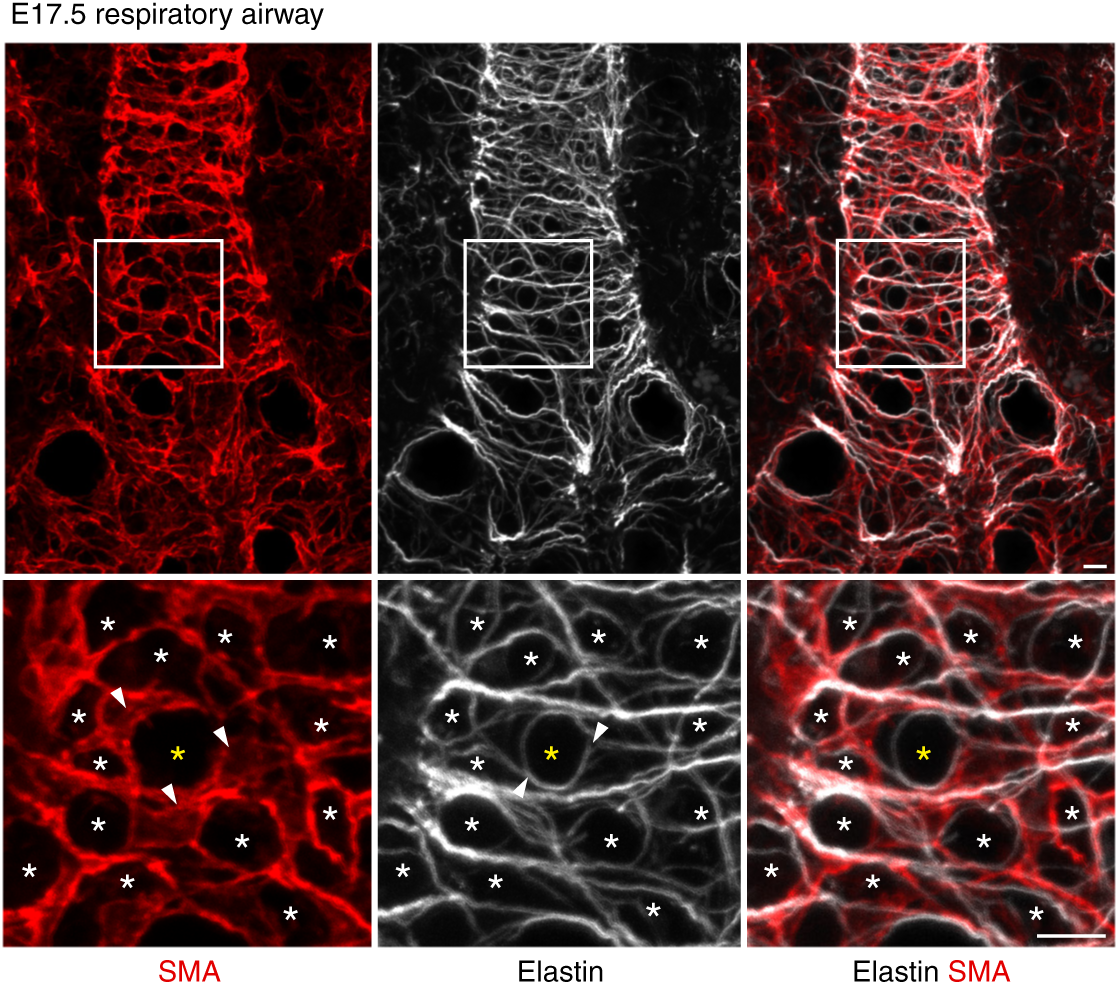
Nascent myofibroblasts align with elastic fibers on respiratory airways. Confocal projection with close-up of boxed region showing myofibroblasts (red) organized as rings (asterisks) and aligned with elastic fibers (white) on the stalk of a respiratory airway in an E17.5 wild-type lung stained for SMA (smooth muscle actin, red) and elastin (detected by hydrazide; white). Note that several myofibroblasts (arrowheads in red panel) and fibers (arrowheads in white panel) form a ring (yellow asterisk). Scale bars, 50 µm.

**Figure S9.**
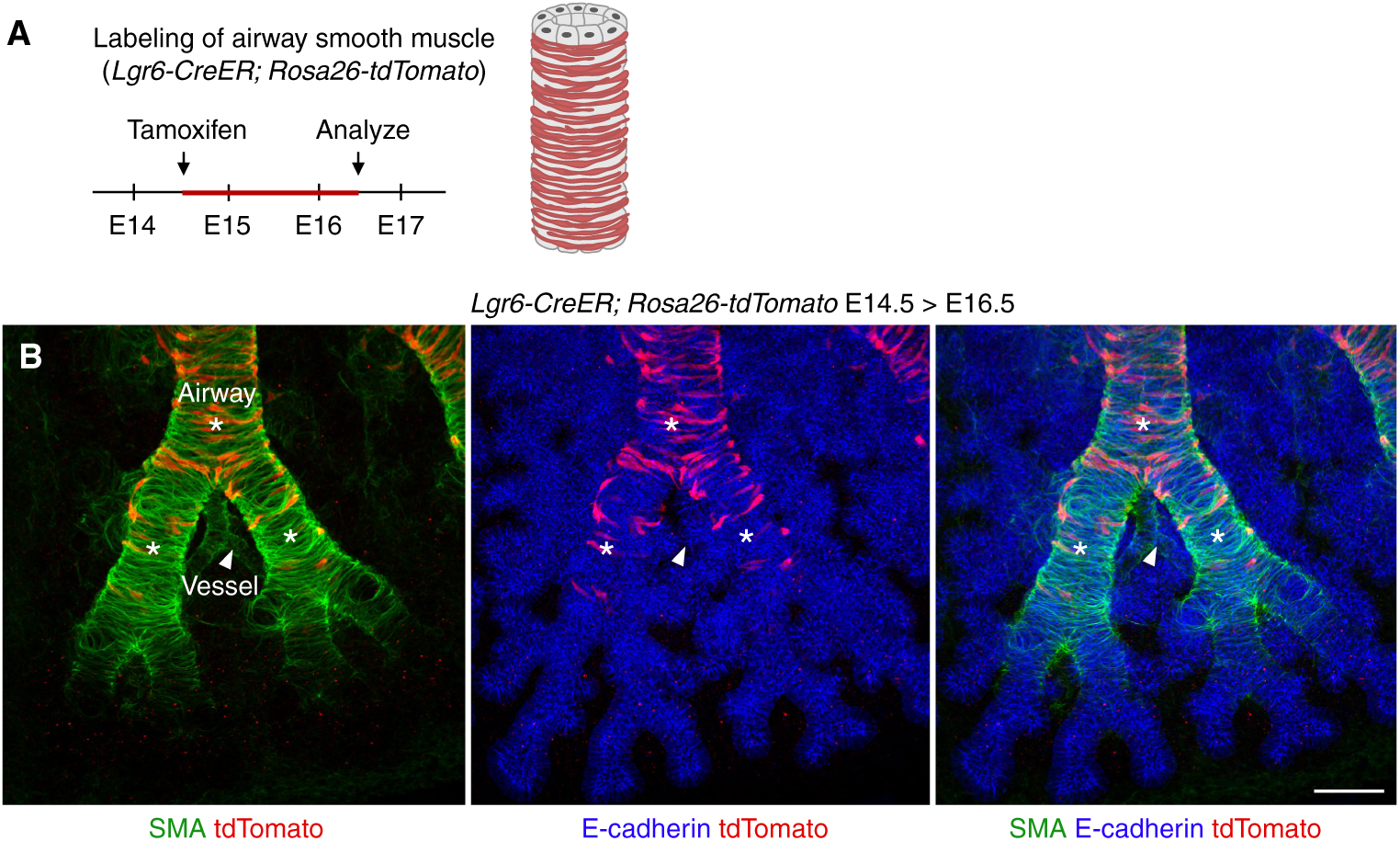
Strategy to label airway smooth muscle. (**A**) Diagram showing genetic strategy to label smooth muscle on embryonic airways. *Lgr6-CreER; Rosa26-tdTomato* lungs were dosed at E14.5 with saturating (4 mg) tamoxifen to label smooth muscle on airway stalks with heritable tdTomato expression (red bar on diagram) and the labeling was analyzed at E16.5. (**B**) Confocal projections of an E16.5 *Lgr6-CreER; Rosa26-tdTomato* lung stained for SMA (smooth muscle, green), the lineage tag (tdTomato, red) and E-cadherin (epithelium, blue) showing lineage-labeled cells (red) on airways (asterisks), but not vessels (arrowhead). Scale bar, 50 µm.

**Figure S10.**
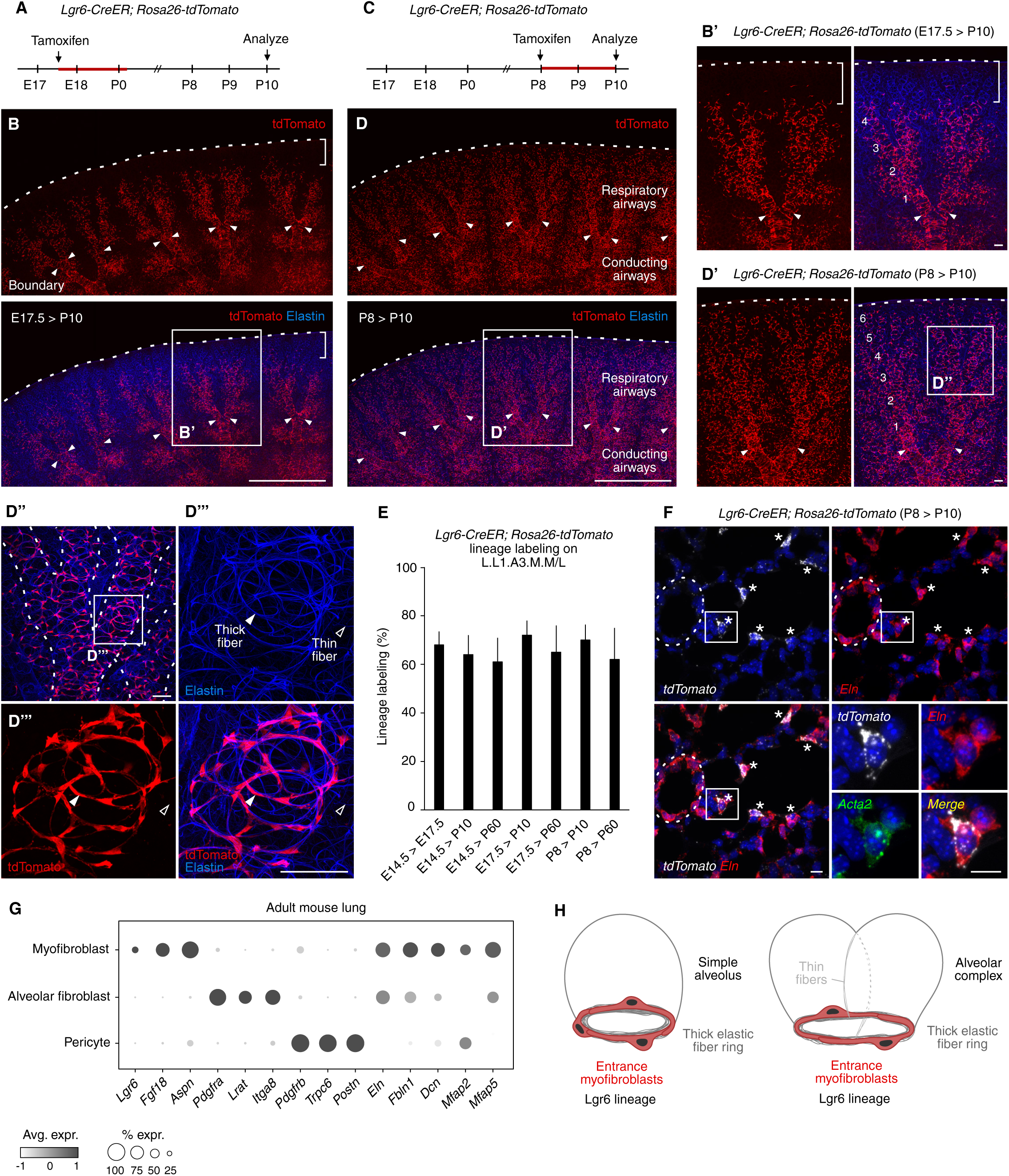
The Lgr6 lineage gives rise to alveolar entrance myofibroblasts. (**A**, **C**) Strategy of *Lgr6-CreER* lineage tracing. *Lgr6-CreER; Rosa26-tdTomato* lungs were dosed at E17.5 (**A**) or P8 (**C**) with saturating (4 mg) tamoxifen to label *Lgr6*-expressing cells with heritable tdTomato expression (red bar on diagram, tamoxifen activity window, with tamoxifen assumed active for 48 hours) and the labeling was analyzed at P10. (**B**, **D**) Confocal projections showing lineage labeling of 3-4 generations (**B’**) and 5-6 generations (**D’-D’’’**) of respiratory airways (numbered) in P10 *Lgr6-CreER; Rosa26-tdTomato* lungs dosed at E17.5 (**B**, **B’**) or at P8 (**D**, **D’-D’’’**). Arrowheads, bronchoalveolar duct junctions. Scale bars, 1 mm (**B**, **D**), 50 µm (**B’**, **D’-D’’’**). (**D’’’**) Lgr6 lineage-labeled cells are aligned with thick (filled arrowhead) but not thin (open arrowhead) elastic fibers. (**E**) Quantification of the efficiency of labeling using *Lgr6-CreER; Rosa26-tdTomato* (data shown as mean ± s.d., n=200 cells scored on L.L1.A3.M.M/L respiratory airways, n=3 mice per group). (**F**) Single-molecule in situ hybridization for *tdTomato* (white), *Eln* (tropoelastin, red) and *Acta2* (SMA, green) in P10 *Lgr6-CreER; Rosa26-tdTomato* lung labeled at P8. Lgr6 lineage-labeled cells (asterisks) express *Eln* transcripts. Dashed oval, vessel wall. Scale bars, 10 µm. (**G**) Dot plot showing expression of tropoelastin (*Eln*) and matrix molecules that associate with elastic fibers (*Fbln1*, fibulin-1; *Dcn*, decorin; *Mfap2* and *Mfap5*, microfibril-associated glycoproteins) in entrance myofibroblasts (identified by expression of *Lgr6*, *Fgf18*, *Aspn*) and alveolar fibroblasts (*Pdgfra*, *Lrat*, *Itga8*), but not pericytes (*Pdgfrb*, *Trpc6*, *Postn*). The cell types were annotated in Tabula Muris Senis scRNAseq Smart-seq2 data for adult mouse lung (*63*). (**H**) Schematic of entrance myofibroblasts in simple (left) and complex (right) alveoli.

**Figure S11.**
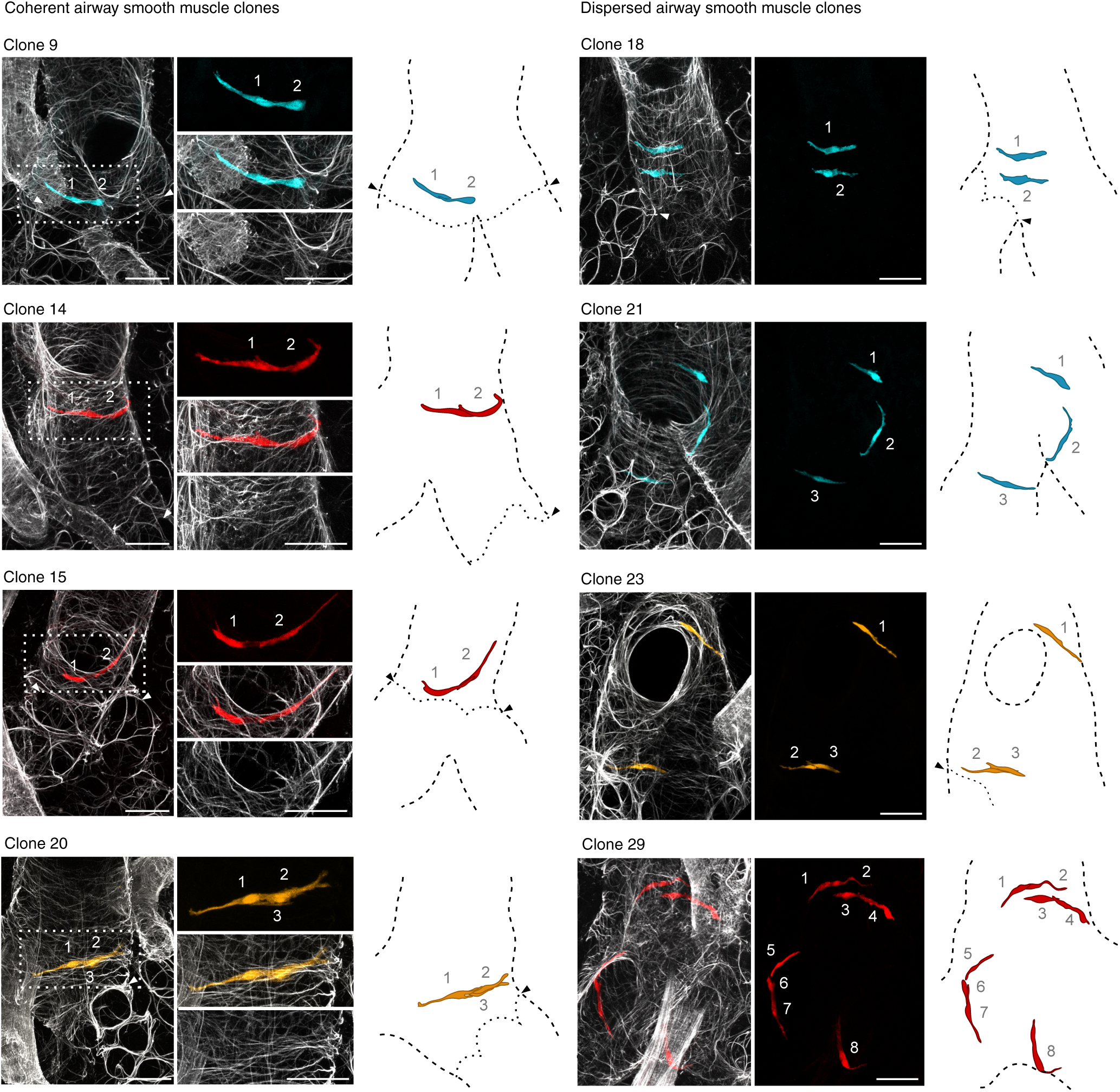
Airway clones generated by tamoxifen induction of *SMMHC-CreER; Rosa26-Rainbow*. Confocal projections (left) and diagrams (right) of airway smooth muscle clones (left panel shows coherent clones, right panel dispersed clones) generated by injecting pregnant *SMMHC-CreER; Rosa26-Rainbow* females with a limiting dose (0.05 mg) of tamoxifen at E15 to induce rare recombination events. Endogenous Cerulean, mCherry, and mOrange fluorescence was analyzed on postnatal day 10 in vibratome sections stained with fluorescent hydrazide (elastin, white) and cleared with Cubic1. Clones with vascular labeling were not analyzed. Cells in each clone are numbered. See also Table S2 for a complete list of clones. Scale bars, 50 µm.

**Figure S12.**
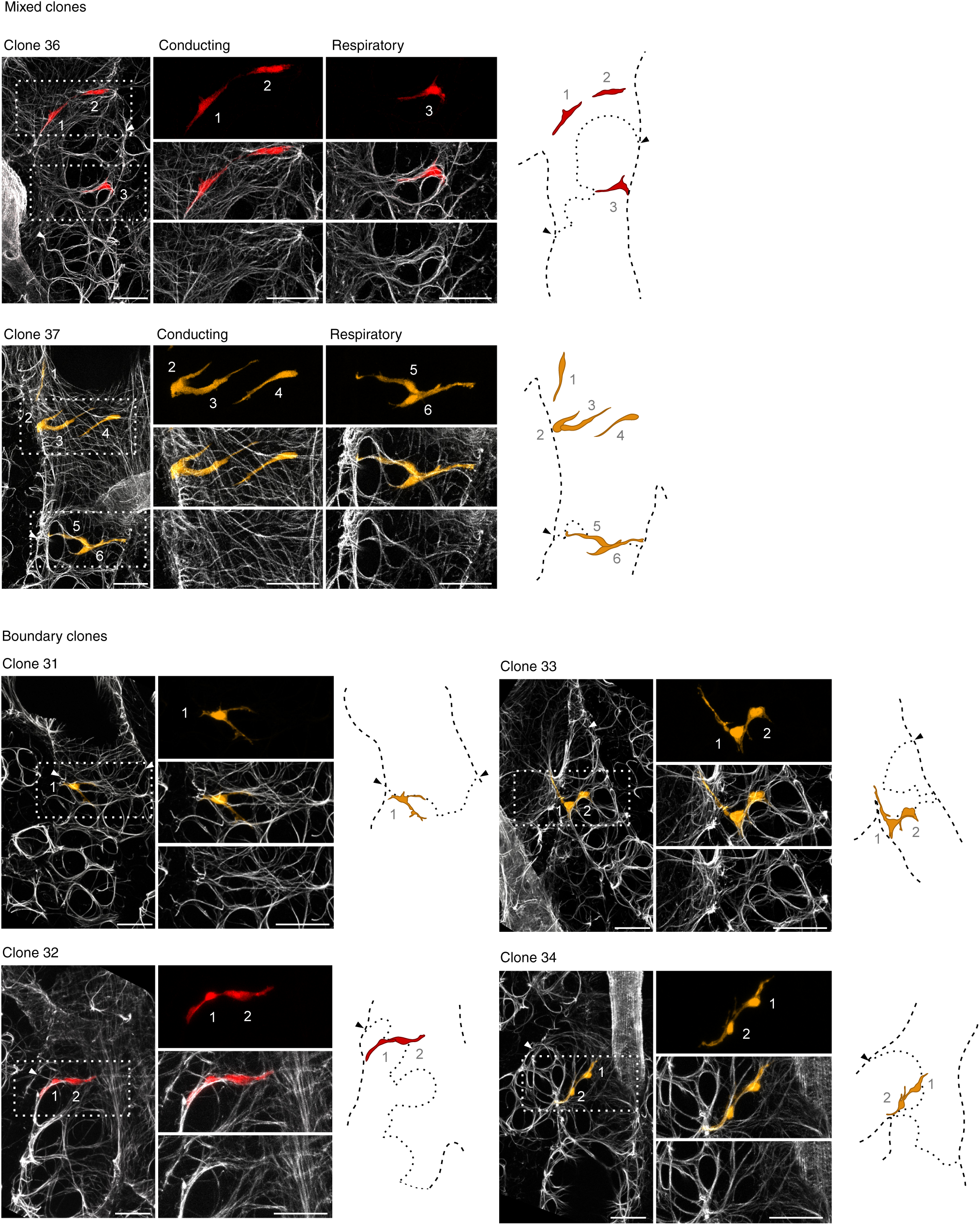
Mixed and boundary clones generated by tamoxifen induction of *SMMHC-CreER; Rosa26-Rainbow*. Confocal projections (left) and diagrams (right) of mixed clones spanning conducting and respiratory airways (top) or clones at the boundary between conducting and respiratory airways (bottom) on postnatal day 10 generated by injecting pregnant *SMMHC-CreER; Rosa26-Rainbow* females with a limiting dose (0.05 mg) of tamoxifen at E15 to induce rare recombination events. The clones are composed of elongated cells with circumferential orientation around airway stalks (airway smooth muscle) and spindle-shaped cells aligned with elastic fibers at the alveolar entrance (myofibroblasts). Cells in each single-color clone are numbered with close-ups of boxed regions at right showing split channels. See also Table S2 for a complete list of clones. Scale bars, 50 µm.

**Figure S13.**
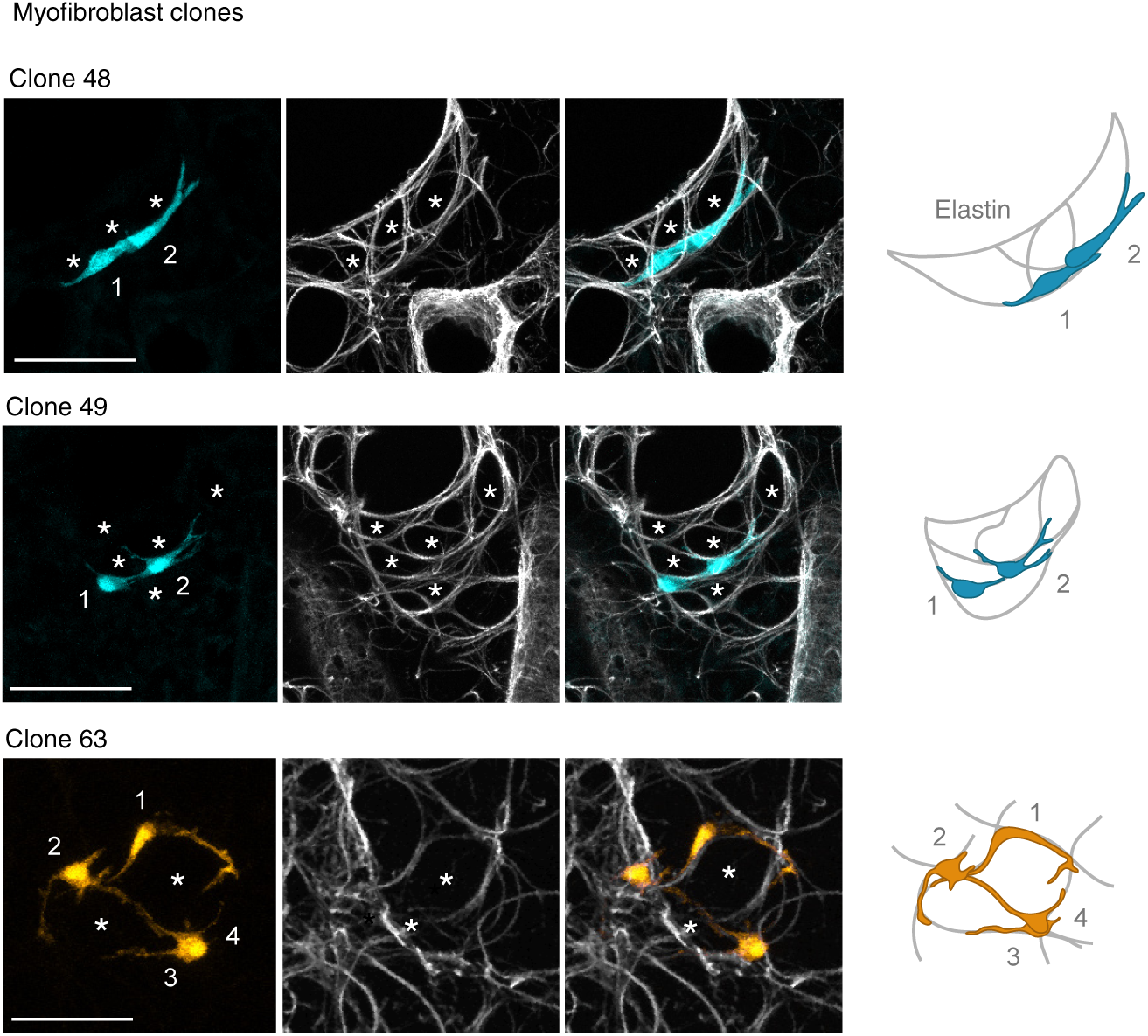
Myofibroblast clones generated by tamoxifen induction of *SMMHC-CreER; Rosa26-Rainbow*. Confocal projections (left) and diagrams (right) as in Fig. S12 showing pure myofibroblast clones on postnatal day 10 generated by injecting pregnant *SMMHC-CreER; Rosa26-Rainbow* females with a limiting dose (0.05 mg) of tamoxifen at E15 to induce rare recombination events. The clones are composed of cells located exclusively on respiratory airways. Cells in each clone are numbered. Asterisks denote alveolar lumens. Note that individual alveolar entrance rings are incompletely labeled suggesting they do not arise by clonal proliferation. See also Table S2 for a complete list of clones. Scale bars, 50 µm.

**Figure S14.**
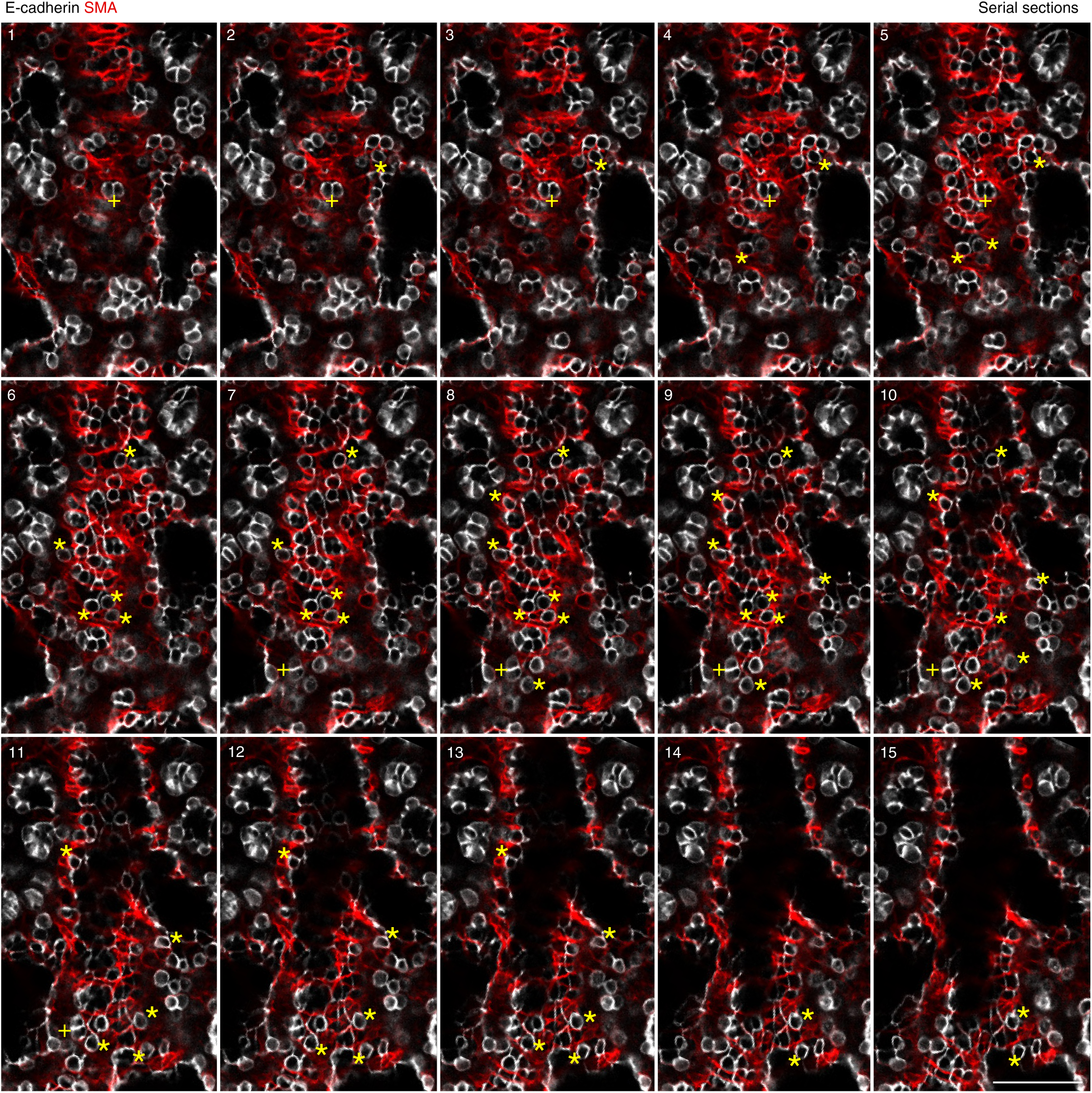
Epithelial pattern and budding on airway stalks through the nascent myofibroblasts. Series of confocal sections (corresponding to the projection image shown in Fig. 5A) showing that single cuboidal epithelial cells (yellow asterisks), or small groups of 2-3 cells (yellow crosses), are selected and protrude through smooth muscle (red) on airway stalks of an E17.5 wild-type lung stained for SMA (smooth muscle actin, red) and E-cadherin (epithelium, white). Scale bar, 50 µm.

**Figure S15.**
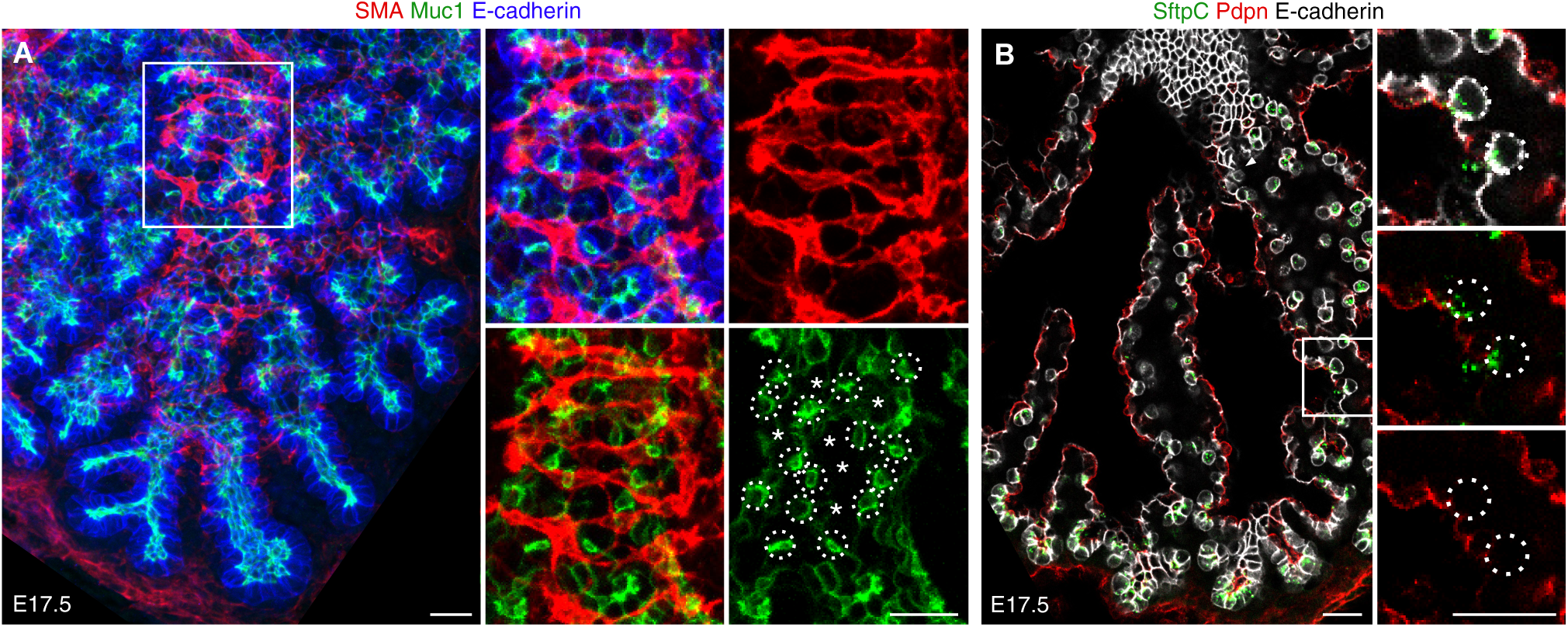
The budding single cuboidal epithelial cells are nascent AT2 cells. (**A**) Confocal projection of an E17.5 wild-type lung stained for SMA (smooth muscle actin, red), Muc1 (progenitors and AT2 cells, green), and E-cadherin (epithelium, blue) with close-up of boxed region showing that single cuboidal epithelial cells (dotted circles) express Muc1 (green), whereas neighboring cells (asterisks; partially flattened AT1 cells) are Muc1-negative/low. Scale bars, 20 µm. (**B**) Single confocal slice of an E17.5 wild-type lung stained for SftpC (progenitors and AT2 cells, green), Pdpn (progenitors and AT1 cells, red) and E-cadherin (epithelium, white). Close-up of boxed region shows that the single cuboidal epithelial cells (dotted circles) are nascent AT2 cells that express SftpC, but not Pdpn. Cuboidal cells in distal tips co-express SftpC and Pdpn. They are progenitors that have not yet undergone differentiation. Scale bars, 20 µm.

**Figure S16.**
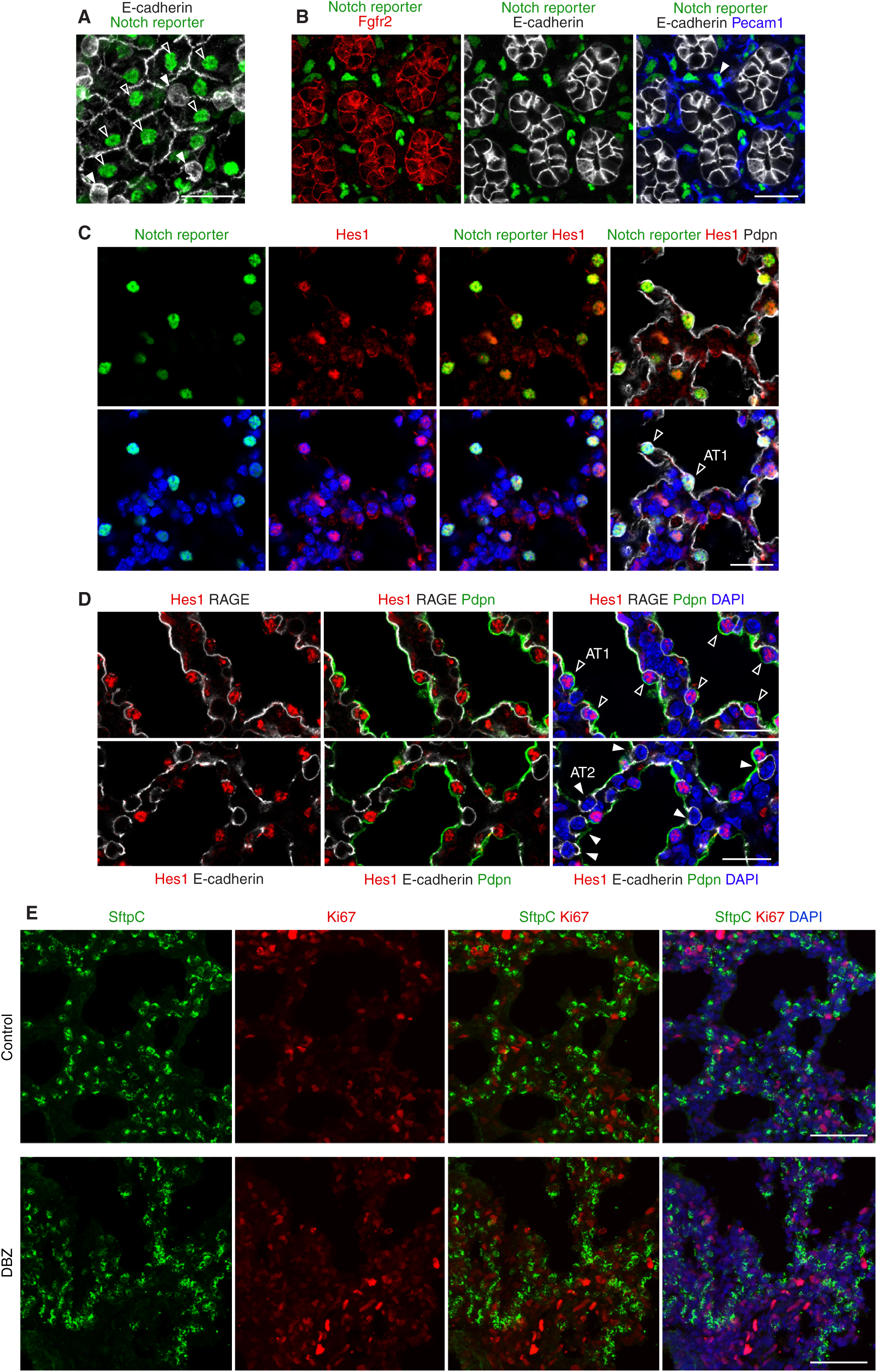
The Notch signaling pathway is activated in nascent AT1 cells. (**A**) Confocal projection of an E18.5 *CBF1-H2B-Venus* lung immunostained for GFP to detect nuclear Venus (green) and for E-cadherin (epithelium, white). Note Notch reporter activity (green) in nascent AT1 cells (open arrowheads) but not nascent AT2 cells (solid arrowheads). Scale bar, 20 µm. (**B**) Confocal optical section of an E16.5 *CBF1-H2B-Venus* (Notch reporter) lung immunostained for GFP to detect nuclear Venus (green), Pecam1 (blue), E-cadherin (epithelium, white) and Fgfr2 (red). Note Notch activity (green) in endothelial plexus (blue), but not epithelial progenitors (white) that express Fgfr2 (red). Scale bar, 20 µm. (**C**) Confocal optical section of an E18.5 *CBF1-H2B-Venus* lung immunostained for GFP to detect Notch reporter (nuclear Venus, green), the Notch-induced gene Hes1 (red), Pdpn (progenitors and AT1 cells, white), and DAPI (nuclei, blue). Note colocalization of Hes1 (red) with nuclear Venus (green) in nascent AT1 cells (open arrowheads). Scale bar, 20 µm. (**D**) Confocal optical slides of an E18.5 wild-type lung immunostained for Hes1 (red), RAGE (AT1 cells, top panel, white) or E-cadherin (epithelium, bottom panel, white), Pdpn (green; progenitor and AT1 marker with apical localization), and DAPI (nuclei, blue). Note Hes1 localizes to nuclei of nascent AT1 cells (open arrowheads), but not AT2 cells (solid arrowheads). Scale bars, 20 µm. (**E**) Analysis of AT2 cell proliferation in embryonic lungs treated with Notch inhibitor DBZ. Confocal projection images showing alveolar epithelium of vehicle (control, top) or DBZ-treated (bottom) E18.5 lungs immunostained for SftpC (progenitors and AT2 cells, green), proliferation marker Ki67 (red) and DAPI (blue). DBZ (30 µM per kilogram body weight) or vehicle were injected daily in wild-type mice (E16-E18). See Fig. 6E for quantification of proliferation. Scale bars, 50 µm.

**Figure S17.**
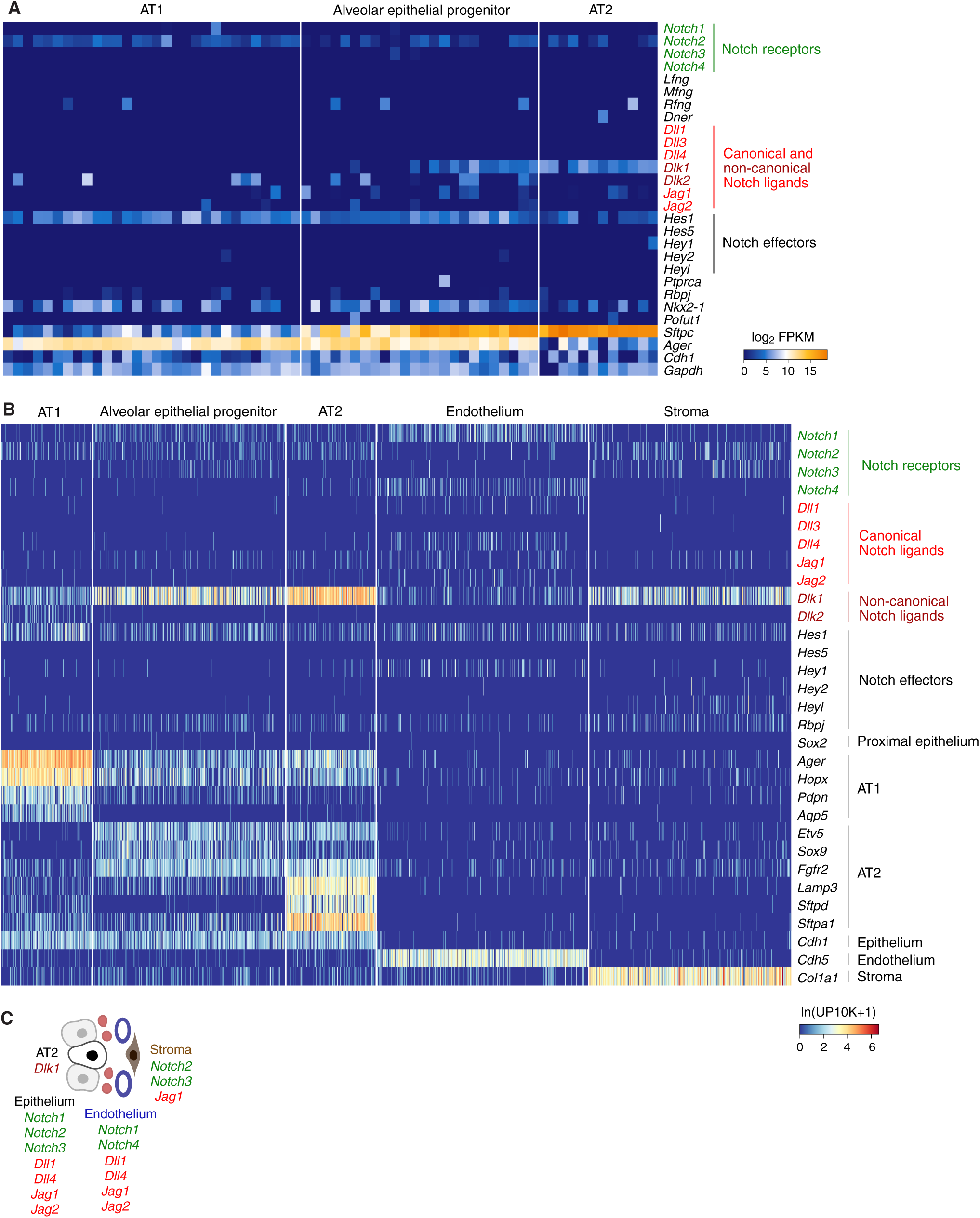
Expression of Notch receptors and ligands in developing alveolar epithelial, endothelial, and stromal cells. (**A**) Heatmap showing normalized expression levels in individual alveolar epithelial cells from E18.5 mouse lung (scRNAseq data from Treutlein et al. (*47*)) for Notch receptors (green), ligands (red), and effectors. FPKM, fragments per kilobase of transcript per million mapped reads; *Sftpc*, progenitor and AT2 marker; *Ager* (RAGE), progenitor and AT1 marker; *Cdh1* (E-cadherin), epithelial marker; *Gapdh,* ubiquitous control. (B) Heatmap showing normalized expression levels in individual alveolar epithelial, endothelial, and stromal cells from developing mouse lung (E16.5 for epithelium, endothelium, and stroma; E18.5 for epithelium; scRNAseq data from Cohen et al. (*74*)) for Notch receptors (green), ligands (red), and effectors. UP10K, unique molecular identifiers per ten thousand. (**C**) Diagram summarizing expression of Notch receptors and showing potential cellular sources of the Notch signal in nascent alveolar buds. Multiple Notch receptors (green), canonical ligands (red), and non-canonical ligands (dark red) are expressed by developing alveolar epithelial cells (progenitors, AT2 and AT1 cells). The phenotype observed upon Notch inhibition in cultured alveolar epithelial progenitors (see Fig. 6F) suggests lateral inhibition of AT2 fate. The expression of *Dlk1* in nascent AT2 cells is also consistent with a potential cis-inhibitory activity of Dlk1 to repress Notch signaling in AT2 cells (Dlk1 lacks the critical Notch binding (DSL) domain) (*98*). Alveolar endothelial and stromal cells are additional potential sources of a Notch signal.

**Table S1.** Cre and CreER drivers used for clonal labeling and lineage tracing.

| **Cre/CreER driver** | **Design** | **Recombination (embryonic)** | **Recombination (E17.5-P10)** | **Recombination (adult)** | **Reference** |
| --- | --- | --- | --- | --- | --- |
| *Shh-GFP-Cre* | knock-in | Epithelial cells | Epithelial cells including AT1/2 cells | Epithelial cells including AT1/2 cells | (82) |
| *Foxa2-CreER* | knock-in | Epithelial cells | Epithelial cells including AT1/2 cells | Epithelial cells including AT1/2 cells | (81) |
| *Hopx-CreER* | knock-in | n.d. | n.d. | AT1 cells, aerocytes | (80) |
| *VE-cadherin-CreER* | transgene | Endothelial cells | Endothelial cells | Endothelial cells | (83) |
| *Apelin-CreER* | knock-in | Plexus | Plexus, emerging aerocytes | Aerocytes | (84) |
| *Aplnr-CreER* | transgene | Plexus | Plexus, emerging general capillary cells | General capillary cells | (85) |
| *Notch3-CreER* | knock-in | n.d. | n.d. | VSM and alveolar pericytes | (88) |
| *Pdgfra-CreER* | transgene | n.d. | n.d. | Stromal cells including alveolar pericytes | (86) |
| *Pdgfra-CreER* | knock-in | n.d. | n.d. | Stromal cells including alveolar fibroblasts | (87) |
| *Gli1-CreER* | knock-in | n.d. | n.d. | Stromal cells including alveolar fibroblasts | (89) |
| *Lgr6-EGFP-IRES-CreERT2* | knock-in | ASM | ASM, entrance myofibroblasts | ASM, entrance myofibroblasts | (60) |
| *SMA-CreER* | transgene | ASM, VSM | ASM, VSM, myofibroblasts | ASM, VSM | (90) |
| *SMMHC-CreER* | transgene | ASM, VSM | ASM, VSM, myofibroblasts | ASM, VSM | (64) |

ASM, airway smooth muscle; VSM, vascular smooth muscle

**Table S2.** Airway clones generated by tamoxifen induction of SMMHC-CreER; Rosa26-Rainbow.

| Clone No. | No. cells | Position | Cell Type | Arrangement | Color |
| --- | --- | --- | --- | --- | --- |
| 1 | 1 | Conducting | Airway smooth muscle | n/a | Cer |
| 2 | 1 | Conducting | Airway smooth muscle | n/a | Cer |
| 3 | 1 | Conducting | Airway smooth muscle | n/a | Cer |
| 4 | 1 | Conducting | Airway smooth muscle | n/a | Cer |
| 5 | 1 | Conducting | Airway smooth muscle | n/a | mOr |
| 6 | 1 | Conducting | Airway smooth muscle | n/a | mOr |
| 7 | 1 | Conducting | Airway smooth muscle | n/a | mOr |
| 8 | 1 | Conducting | Airway smooth muscle | n/a | mCh |
| 9 | 2 | Conducting | Airway smooth muscle | Coherent | Cer |
| 10 | 2 | Conducting | Airway smooth muscle | Coherent | Cer |
| 11 | 2 | Conducting | Airway smooth muscle | Coherent | mOr |
| 12 | 2 | Conducting | Airway smooth muscle | Coherent | mOr |
| 13 | 2 | Conducting | Airway smooth muscle | Coherent | mOr |
| 14 | 2 | Conducting | Airway smooth muscle | Coherent | mCh |
| 15 | 2 | Conducting | Airway smooth muscle | Coherent | mCh |
| 16 | 2 | Conducting | Airway smooth muscle | Coherent | mCh |
| 17 | 2 | Conducting | Airway smooth muscle | Coherent | mCh |
| 18 | 2 | Conducting | Airway smooth muscle | Dispersed | Cer |
| 19 | 2 | Conducting | Airway smooth muscle | Dispersed | Cer |
| 20 | 3 | Conducting | Airway smooth muscle | Coherent | mOr |
| 21 | 3 | Conducting | Airway smooth muscle | Dispersed | Cer |
| 22 | 3 | Conducting | Airway smooth muscle | Dispersed | Cer |
| 23 | 3 | Conducting | Airway smooth muscle | Dispersed | mOr |
| 24 | 4 | Conducting | Airway smooth muscle | Coherent | mCh |
| 25 | 4 | Conducting | Airway smooth muscle | Dispersed | mOr |
| 26 | 4 | Conducting | Airway smooth muscle | Dispersed | mOr |
| 27 | 4 | Conducting | Airway smooth muscle | Dispersed | mCh |
| 28 | 5 | Conducting | Airway smooth muscle | Dispersed | Cer |
| 29 | 8 | Conducting | Airway smooth muscle | Dispersed | mCh |
| 30 | 13 | Conducting | Airway smooth muscle | Dispersed | mCh |
| 31 | 1 | Boundary | Hybrid | n/a | mOr |
| 32 | 2 | Boundary | Hybrid | Coherent | mCh |
| 33 | 2 | Boundary | Hybrid | Coherent | mOr |
| 34 | 2 | Boundary | Hybrid | Coherent | mOr |
| 35 | 2 | Conducting+respiratory | Mixed | Coherent | mCh |
| 36 | 3 | Conducting+respiratory | Mixed | Dispersed | mCh |
| 37 | 6 | Conducting+respiratory | Mixed | Dispersed | mOr |
| 38 | 10 | Conducting+respiratory | Mixed | Dispersed | Cer |
| 39 | 1 | Respiratory | Myofibroblast | n/a | Cer |
| 40 | 1 | Respiratory | Myofibroblast | n/a | Cer |
| 41 | 1 | Respiratory | Myofibroblast | n/a | Cer |
| 42 | 1 | Respiratory | Myofibroblast | n/a | Cer |
| 43 | 1 | Respiratory | Myofibroblast | n/a | mOr |
| 44 | 1 | Respiratory | Myofibroblast | n/a | mOr |
| 45 | 1 | Respiratory | Myofibroblast | n/a | mOr |
| 46 | 1 | Respiratory | Myofibroblast | n/a | mCh |
| 47 | 1 | Respiratory | Myofibroblast | n/a | mCh |
| 48 | 2 | Respiratory | Myofibroblast | Coherent | Cer |
| 49 | 2 | Respiratory | Myofibroblast | Coherent | Cer |
| 50 | 2 | Respiratory | Myofibroblast | Coherent | Cer |
| 51 | 2 | Respiratory | Myofibroblast | Coherent | Cer |
| 52 | 2 | Respiratory | Myofibroblast | Coherent | mOr |
| 53 | 2 | Respiratory | Myofibroblast | Coherent | mOr |
| 54 | 2 | Respiratory | Myofibroblast | Coherent | mOr |
| 55 | 2 | Respiratory | Myofibroblast | Coherent | mOr |
| 56 | 2 | Respiratory | Myofibroblast | Coherent | mOr |
| 57 | 2 | Respiratory | Myofibroblast | Coherent | mOr |
| 58 | 2 | Respiratory | Myofibroblast | Coherent | mCh |
| 59 | 2 | Respiratory | Myofibroblast | Coherent | mCh |
| 60 | 2 | Respiratory | Myofibroblast | Coherent | mCh |
| 61 | 3 | Respiratory | Myofibroblast | Coherent | mOr |
| 62 | 3 | Respiratory | Myofibroblast | Dispersed | mOr |
| 63 | 4 | Respiratory | Myofibroblast | Coherent | mOr |
| 64 | 4 | Respiratory | Myofibroblast | Dispersed | mCh |
| 65 | 8 | Respiratory | Myofibroblast | Dispersed | mCh |
| 66 | 12 | Respiratory | Myofibroblast | Dispersed | mOr |
| 67 | 14 | Respiratory | Myofibroblast | Dispersed | mCh |
| 68 | 17 | Respiratory | Myofibroblast | Dispersed | mCh |

